# GtcA is required for LTA glycosylation in *Listeria monocytogenes* serovar 1/2a and *Bacillus subtilis*

**DOI:** 10.1101/2019.12.12.873851

**Authors:** Jeanine Rismondo, Talal F. M. Haddad, Yang Shen, Martin J. Loessner, Angelika Gründling

**Author notes:** To whom correspondence should be addressed: Angelika Gründling –.

## Abstract

The cell wall polymers wall teichoic acid (WTA) and lipoteichoic acid (LTA) are often modified with glycosyl and D-alanine residues. Recent studies have shown that a three-component glycosylation system is used for the modification of LTA in several Gram-positive bacteria including *Bacillus subtilis* and *Listeria monocytogenes*. In the *L. monocytogenes* 1/2a strain 10403S, the cytoplasmic glycosyltransferase GtlA is thought to use UDP-galactose to produce the C_55_-P-galactose lipid intermediate, which is transported across the membrane by an unknown flippase. Next, the galactose residue is transferred onto the LTA backbone on the outside of the cell by the glycosyltransferase GtlB. Here we show that GtcA is necessary for the glycosylation of LTA in *L. monocytogenes* 10403S and *B. subtilis* 168 and we hypothesize that these proteins act as C_55_-P-sugar flippases. With this we revealed that GtcA is involved in the glycosylation of both teichoic acid polymers in *L. monocytogenes* 10403S, namely WTA with N-acetylglucosamine and LTA with galactose residues. These findings indicate that the *L. monocytogenes* GtcA protein can act on different C_55_-P-sugar intermediates. Further characterization of GtcA in *L. monocytogenes* led to the identification of residues essential for its overall function as well as residues, which predominately impact WTA or LTA glycosylation.

**GRAPHICAL ABSTRACT:** 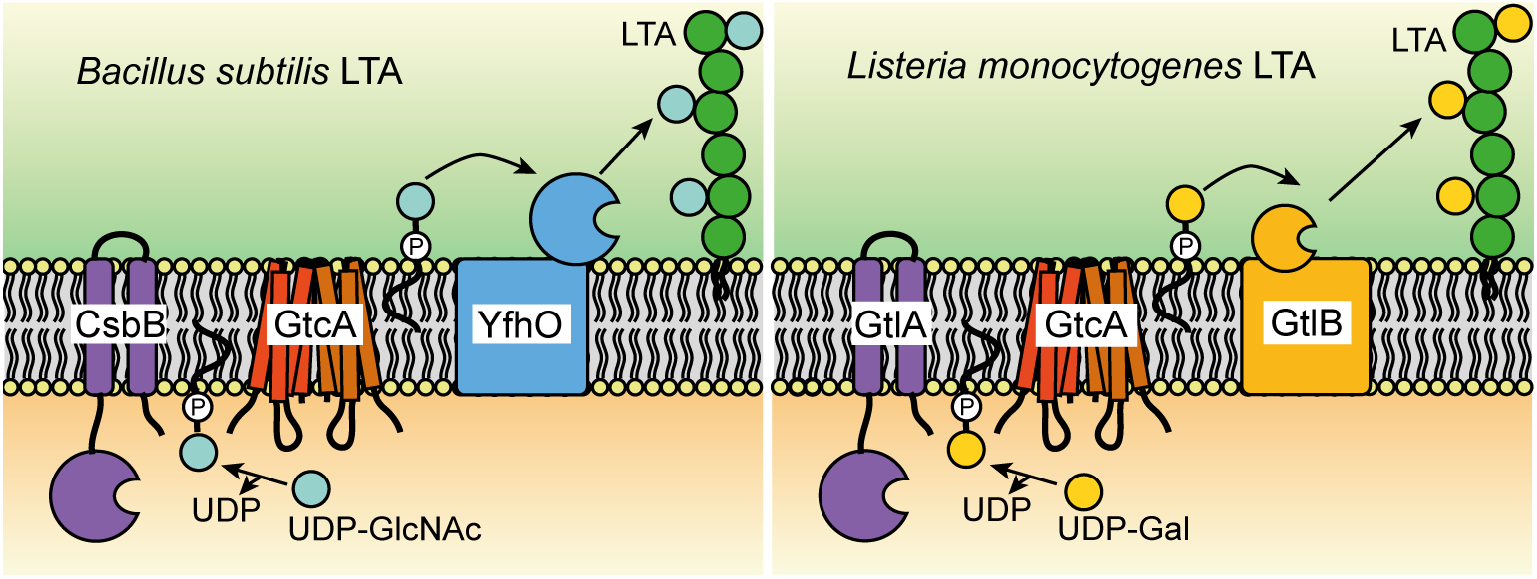

## 1. INTRODUCTION

Bactoprenol, also referred to as undecaprenyl phosphate (C_55_-P), is used for a wide range of processes in bacteria including the biosynthesis of peptidoglycan, lipopolysaccharides (LPS) and teichoic acids (Harkness and Braun, 1989, Kennedy, 1987, Weissborn et al., 1991, Soldo et al., 2002). This molecule is an essential lipid-carrier to which sugars, monomeric subunits of cell wall components or complete polymers are linked in the cytoplasm of the cell. The lipid-linked intermediates are subsequently moved across the membrane by flippase enzymes or transporters and finally incorporated on the outside of the cell in diverse cell wall structures. However, the mechanism by which the different lipid-linked intermediates are flipped or transported across the membrane as well as the proteins involved in such processes are not well characterized.

Teichoic acids are important cell wall polymers produced by Gram-positive bacteria. They are either covalently linked to the peptidoglycan and referred to as wall teichoic acid (WTA) or embedded in the cytoplasm via a glycolipid anchor and called lipoteichoic acid (LTA) (Araki and Ito, 1989, Neuhaus and Baddiley, 2003, Percy and Gründling, 2014). These cell wall polymers are essential for the maintenance of the cell integrity and play an important role in antimicrobial resistance, cation homeostasis, regulation of peptidoglycan autolysins, cell division and virulence and the simultaneous absence of both, LTA and WTA, has been shown to be lethal in *Staphylococcus aureus* and *Bacillus subtilis* (Brown et al., 2012, Spears et al., 2016, Schirner et al., 2009, Percy and Gründling, 2014, Brown et al., 2013). The bactoprenol lipid carrier molecular C_55_-P is required at multiple steps during the synthesis and modification process of teichoic acids in Gram-positive bacteria.

In *B. subtilis*, the synthesis of WTA begins in the cytoplasm of the cell with the glycosyltransferase (GT) TagO, which transfers N-acetyl glucosamine (GlcNAc) phosphate from UDP-GlcNAc onto undecaprenyl phosphate (Soldo et al., 2002). Next, N-acetylmannosamine (ManNAc) is attached onto C_55_-PP-GlcNAc by TagA, followed by the transfer of glycerol phosphate (GroP) units by TagB and TagF (Bhavsar et al., 2005, Ginsberg et al., 2006, Schertzer and Brown, 2003, Pereira et al., 2008). Following the synthesis of the WTA polymer, the GroP backbone is further modified with glucose residues on the inside of the cell by the cytoplasmic glycosyltransferase (GT) TagE. The synthesized and glycosylated WTA polymer is subsequently transported across the membrane by the ABC-transporter TagGH (Lazarevic and Karamata, 1995). After the export, WTA is modified with D-Alanine residues by enzymes encoded in the *dlt* operon and linked to the peptidoglycan layer by Lcp enzymes (Kawai et al., 2011, Perego et al., 1995). Other Gram-positive bacteria, including most *S. aureus* strains, *Listeria monocytogenes* strains, and *B. subtilis* strain W23 produce WTA with a ribitolphosphate (RboP) backbone (Brown et al., 2010, Weidenmaier and Peschel, 2008, Uchikawa et al., 1986). The enzymes involved in this process are encoded by the *tar/tag* genes (Weidenmaier and Peschel, 2008, Brown et al., 2010). WTA of *S. aureus* is also modified on the inside of the cell with GlcNAc residues by the cytoplasmic glycosyltransferases (GTs) TarM, TarS and TarP before it is exported and attached to the peptidoglycan layer (Allison et al., 2011, Gerlach et al., 2018, Xia et al., 2010, Brown et al., 2012, Dengler et al., 2012). Recently, a different mechanism has been proposed for the glycosylation of WTA with GlcNAc residues in *L. monocytogenes* serovar 1/2a strains (Rismondo et al., 2018). In this case, the cytoplasmic GT Lmo2550 utilizes UDP-GlcNAc to form C_55_-P-GlcNAc, a lipid linked sugar intermediate, which is flipped across the membrane by the putative flippase GtcA (Lmo2549) (Eugster et al., 2011, Promadej et al., 1999, Cheng et al., 2008). In the next step, the glycosyltransferase YfhO (Lmo1079) is thought to transfer the GlcNAc residue from the C_55_-P-GlcNAc intermediate onto the WTA backbone on the outside of the cell (Eugster et al., 2015, Rismondo et al., 2018)(Fig. 1A).

**Figure 1:**
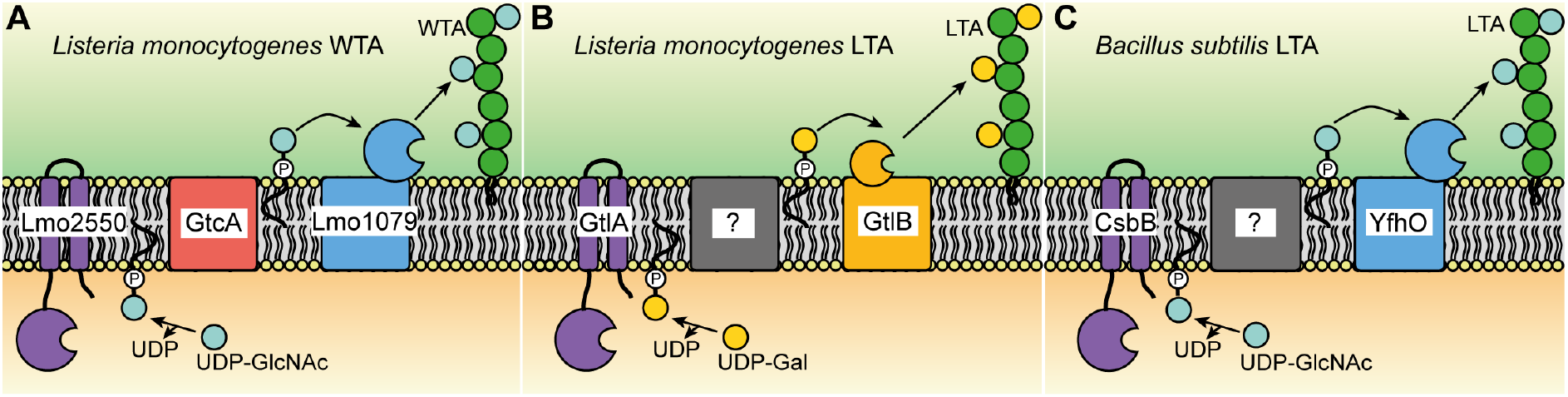
Overview over the proposed WTA and LTA glycosylation process in *L. monocytogenes* serovar 1/2a strains and LTA glycosylation in *B. subtilis*. (A) Model of the WTA glycosylation process in *L. monocytogenes*. The cytoplasmic glycosyltransferase (GT) Lmo2550 uses UDP-GlcNAc to form a C_55_-P-GlcNAc intermediate that is flipped across the membrane by the putative flippase GtcA (Eugster et al., 2011, Promadej et al., 1999, Cheng et al., 2008). Lmo1079 (YfhO), then transfers the GlcNAc residue onto the WTA backbone on the outside of the cell (Eugster et al., 2015). (B) Model of the LTA glycosylation process in *L. monocytogenes*. GtlA, the putative cytoplasmic GT, forms a C_55_-P-galactose intermediate (Percy et al., 2016) that is transported across the membrane by an unknown flippase enzyme. The galactose residue is then transferred onto the LTA backbone by the likely GT enzyme GtlB (Rismondo et al., 2018). (C) Model of the LTA glycosylation process in *B. subtilis*. The cytoplasmic GT CsbB transfers GlcNAc residues to the C_55_-P lipid carrier (Rismondo et al., 2018). The C_55_-P-GlcNAc intermediate is subsequently transported across the membrane by an unknown flippase and the GlcNAc residue is transferred onto the LTA backbone by YfhO on the outside of the cell (Rismondo et al., 2018).

A similar three enzyme glycosylation process has been suggested for the glycosylation of the LTA polymer, which is synthesized on a glycolipid anchor on the outside of the cell and hence can only be glycosylated extracellularly (Iwasaki et al., 1989, Fischer, 1994, Mancuso and Chiu, 1982, Yokoyama et al., 1988) (Fig. 1B and 1C). The synthesis of LTA starts in the cytoplasm with the transfer of two glucose molecules from UDP-glucose to diacylglycerol (DAG) by UgtP and YpfP in *B. subtilis* and *S. aureus*, respectively (Jorasch et al., 1998, Kiriukhin et al., 2001). In *L. monocytogenes*, the glycolipid anchor is produced by LafA and LafB, which transfer a glucose and subsequently a galactose molecule onto DAG (Webb et al., 2009). The glycolipids are subsequently transported across the membrane and this step is in *S. aureus* likely mediated by the membrane protein LtaA (Kiriukhin et al., 2001, Gründling and Schneewind, 2007a). The proteins required for the flipping of the glycolipid have not yet been identified in *B. subtilis* and *L. monocytogenes*. The glycerolphosphate LTA backbone is subsequently polymerized directly on the glycolipid anchor on the outside of the cell by one or multiple lipoteichoic acid synthase (LtaS)-type enzymes (Gründling and Schneewind, 2007b, Webb et al., 2009, Wörmann et al., 2011, Schirner et al., 2009). The LTA GroP backbone is then modified with D-alanine and often additional sugar residues. Biochemical studies led to the suggestion that the LTA glycosylation process starts with the synthesis of a C_55_-P lipid-linked sugar intermediate by a cytoplasmic glycosyltransferase. This intermediate is subsequently flipped across the membrane by a flippase enzyme. In the last step, a glycosyltransferase with extracellular activity transfers the sugar moiety onto the LTA backbone (Fischer, 1994, Iwasaki et al., 1989, Yokoyama et al., 1988, Mancuso and Chiu, 1982). Consistent with this model, a three-component glycosylation system composed of the cytoplasmic glycosyltransferase CsbB, the small membrane protein and putative flippase GtcA and the LTA-specific glycosyltransferase YfhO has recently been shown to be required for the glycosylation of the LTA polymer in *S. aureus* with GlcNAc residues (Kho and Meredith, 2018). In *B. subtilis*, CsbB and YfhO have been identified as likely cytoplasmic and extracellular GTs (Fig. 1C) and mutations in the genes coded for these enzymes led to the absence of glucose residues on LTA (Percy et al., 2016, Rismondo et al., 2018). Here it is interesting to note that as mentioned above the YfhO homolog of *L. monocytogenes*, Lmo1079, is responsible for the glycosylation of WTA with GlcNAc residues and not required for the LTA glycosylation process. This observation raises the question of the sugar- and acceptor-specificity of YfhO-like enzymes, a question that was addressed as part of this study. In *L. monocytogenes* serotype 1/2a strains, *gtlA* and *gtlB* encode the likely cytoplasmic and extracellular GTs involved in the LTA glycosylation process and deletion of these genes leads to the absence of galactose residues on LTA (Fig. 1B) (Percy et al., 2016, Rismondo et al., 2018). For a *L. monocytogenes* serotype 4b strain, it has recently been shown that *gttA*, encoding a membrane-anchored GT with a cytoplasmatically located enzymatic domain, is likely involved in the production of a C_55_-P-galactose intermediate, which is required for the glycosylation of LTA and WTA (Sumrall et al., 2019). However, the proteins involved in the transport of the C_55_-P-sugar intermediates during the LTA glycosylation have not been identified in *B. subtilis* and *L. monocytogenes*. Deletion of *gtcA* in a *L. monocytogenes* serotype 4b strain has been shown to lead to the loss of sugar modifications on WTA (Promadej et al., 1999). While there is evidence in the literature that the absence of GtcA does not impact the structure of LTA in *L. monocytogenes* serovar 4b strains (Promadej et al., 1999), the recent finding that LTA and WTA are glycosylated by similar mechanisms in *L. monocytogenes* 10403S, a serovar 1/2a strain (Rismondo et al., 2018) (Fig. 1A and 1B) prompted us to revisit the involvement of GtcA in the LTA glycosylation process. A GtcA homolog is also present in *B. subtilis*, but its function in the LTA glycosylation process has not been investigated. As part of this study, *gtcA* deletion and complementation strains were constructed in *B. subtilis* 168 and *L. monocytogenes* 10403S and the composition of teichoic acid polymers analyzed using a combination of western-blot, fluorescence microscopy, NMR and mass-spectrometry techniques. Our results show that GtcA is required for the glycosylation of LTA in *B. subtilis* 168 and for LTA and WTA glycosylation in *L. monocytogenes* 10403S, suggesting that the predicted flippase can recognize and transport both C_55_-P-galactose and C_55_-P-GlcNAc intermediates in *L. monocytogenes*. Using a mutagenesis approach, we identified conserved residues that are essential for GtcA protein function, as well as residues that primarily affect LTA or WTA glycosylation in *L. monocytogenes*. To our knowledge, this is the first time that essential residues have been identified in predicted undecaprenyl carrier flippases belonging to the GtrA family of proteins (Allison and Verma, 2000) and this information will help us better understand how members of this large protein family could function as undecaprenyl carrier flippases.

## 2. MATERIALS AND METHODS

### 2.1 Bacterial strains and growth conditions

All strains and plasmids used in this study are listed in Table S1. *Escherichia coli* and *Bacillus subtilis* strains were grown in lysogenic broth (LB) medium and *Listeria monocytogenes* strains in brain heart infusion (BHI) medium at 37°C unless otherwise stated. If necessary, antibiotics and supplements were added to the medium at following concentrations: for *E. coli* cultures, ampicillin (Amp) at 100 μg/ml, chloramphenicol (Cam) at 20 μg/ml and kanamycin (Kan) at 30 μg/ml, for *B. subtilis* cultures, chloramphenicol (Cam) at 5 μg/ml, kanamycin (Kan) at 10 μg/ml, and for *L. monocytogenes* cultures, chloramphenicol (Cam) at 10 μg/ml, kanamycin (Kan) at 30 μg/ml, IPTG at 1 mM.

### 2.2 Strain and plasmid construction

All primers used in this study are listed in Table S2. For the generation of *L. monocytogenes* strains 10403SΔ*gtlA*Δ*gtlB* (or short 10403SΔ*gtlAB*; ANG5195) and 10403SΔ*lmo1079*Δ*lmo2550* (ANG5197), plasmids pKSV7-Δ*gtlB* (ANG4738) and pKSV7-Δ*lmo2550* (ANG2223) were transformed into 10403SΔ*gtlA* (ANG2325) and 10403SΔ*lmo1079* (ANG2794), respectively, and the *gtlB* gene or *lmo2550* gene deleted by allelic exchange using a previously described method (Camilli et al., 1993). The deletion of *gtlB* and *lmo2550* was verified by PCR. For the IPTG-inducible expression of the *csbB-yfhO* operon in *L. monocytogenes*, the *csbB-yfhO* region was amplified using *B. subtilis* 168 chromosomal DNA and primers ANG3143/3144. The resulting PCR product was cut with NcoI and SalI and ligated with plasmid pIMK3 that had been cut with the same enzymes. Plasmid pIMK3-*csbB-yfhO* was recovered in *E. coli* XL1-Blue yielding strain ANG5182. pIMK3-*csbB-yfhO* was subsequently transformed into *L. monocytogenes* strains 10403SΔ*gtlAB* and 10403SΔ*lmo1079*Δ*lmo2550* by electroporation, resulting in the construction of strains 10403SΔ*gtlAB* pIMK3-*csbB-yfhO* (ANG5199) and 10403SΔ*lmo1079*Δ*lmo2550* pIMK3-*csbB-yfhO* (ANG5203).

To generate markerless in-frame deletions in the *L. monocytogenes* genes *gtcA* (*lmo2549*) and *lmo0215*, approximately 1kb-DNA fragments up- and downstream of *gtcA* and *lmo0215* were amplified using primer pairs ANG2979/2980 and ANG2981/2982 (*gtcA*) and ANG3305/3306 and ANG3307/3308 (*lmo0215*). The resulting PCR products were fused through a second PCR using primers ANG2979/2982 (*gtcA*) and ANG3305/3308 (*lmo0215*), cut with BamHI and KpnI and ligated with pKSV7 that had been digested with the same enzymes. Plasmids pKSV7-Δ*gtcA* and pKSV7-Δ*lmo0215* were subsequently recovered in *E. coli* XL1-Blue yielding strains ANG4911 and ANG5619, respectively. Next, the plasmids were electroporated into *L. monocytogenes* 10403S and the *gtcA* and *lmo0215* deleted by allelic exchange yielding strains 10403SΔ*gtcA* (ANG4972) and 10403SΔ*lmo0215* (ANG5638). The deletion of genes *gtcA* and *lmo0215* was verified by PCR. For the construction of a *gtcA* complementation strain, plasmid pIMK3-*gtcA* was constructed enabling the IPTG-dependent expression of *gtcA*. For this purpose, *gtcA* was amplified using primers ANG3036/3037, the resulting PCR product digested with NcoI and SalI and ligated with pIMK3. Plasmid pIMK3-*gtcA* was recovered in *E. coli* XL1-Blue yielding strain ANG5026. Point mutations were introduced into the *gtcA* gene for the expression of GtcA variants with A65S, N69A, V73A, F74A, F91A, R95A, K121A and N132A amino acid substitutions. To this end, primer pairs ANG3036/3216 (A65S), ANG3036/3218 (N69A), ANG3036/3220 (V73A), ANG3036/3222 (F74A), ANG3036/3224 (F91A), ANG3036/3226 (R95A), ANG3036/3228 (K121A) and ANG33036/3230 (N132A) were used to amplify the 5’ end of *gtcA* introducing the appropriate base change(s) as part of the reverse primer sequence. The 3’ end of *<u>gtcA</u>* was amplified using primers ANG3037/3215 (A65S), ANG3037/3217 (N69A), ANG3037/3219 (V73A), ANG3037/3221 (F74A), ANG3037/3223 (F91A), ANG3037/3225 (R95A), ANG3037/3227 (K121A) and ANG3037/3229 (N132A) introducing the appropriate base change(s) as part of the forward primer. The corresponding 5’ and 3’ *gtcA* fragments were fused in a second PCR using primers ANG3036/3037. The resulting PCR products were digested with NcoI and SalI and ligated with pIMK3. The resulting plasmids were recovered in *E. coli* XL1-Blue yielding strains XL1-Blue pIMK3-*gtcA_A65S_* (ANG5620), XL1-Blue pIMK3-*gtcA_N69A_* (ANG5621), XL1-Blue pIMK3-*gtcA_V73A_* (ANG5622), XL1-Blue pIMK3-*gtcA_F74A_* (ANG5623), XL1-Blue pIMK3-*gtcA_F91A_* (ANG5624), XL1-Blue pIMK3-*gtcA_R95A_* (ANG5625), XL1-Blue pIMK3-*gtcA_K121A_* (ANG5626) and XL1-Blue pIMK3-*gtcA_N132A_* (ANG5627). Additionally, pIMK3-*His-gtcA* and derivatives carrying the above described point mutations were constructed allowing for the expression of N-terminally His-tagged GtcA proteins and detection by western blot. *gtcA* was amplified using primers ANG3345/3037. The resulting PCR product was used in a second PCR using primers ANG3346/3037 to attach the sequence of the N-terminal His-tag. The *His-gtcA* fragment was subsequently cut with NcoI and SalI and ligated with pIMK3. Plasmid pIMK3-*His-gtcA* was recovered in *E. coli* XL1-Blue yielding strain ANG5628. For the introduction of the point mutations, primer pairs ANG3345/3216 (A65S), ANG3345/3218 (N69A), ANG3345/3220 (V73A), ANG3345/3222 (F74A), ANG3345/3224 (F91A), ANG3345/3226 (R95A), ANG3345/3228 (K121A) and ANG3345/3230 (N132A) were used to amplify the 5’ end of *gtcA*. The 3’ end of **gtcA** was amplified using primers ANG3037/3215 (A65S), ANG3037/3217 (N69A), ANG3037/3219 (V73A), ANG3037/3221 (F74A), ANG3037/3223 (F91A), ANG3037/3225 (R95A), ANG3037/3227 (K121A) and ANG3037/3229 (N132A). The appropriate fragments were fused in a second PCR using primers ANG3346/3037. The PCR products were cut with NcoI and SalI and fused with pIMK3 that had been cut with the same enzymes. The resulting plasmids were recovered in *E. coli* XL1-Blue yielding strains XL1-Blue pIMK3-*His-gtcA_A65S_* (ANG5629), XL1-Blue pIMK3-*His-gtcA_N69A_* (ANG5630), XL1-Blue pIMK3-*His-gtcA_V73A_* (ANG5631), XL1-Blue pIMK3-*His-gtcA_F74A_* (ANG5632), XL1-Blue pIMK3-*His-gtcA_F91A_* (ANG5633), XL1-Blue pIMK3-*His-gtcA_R95A_* (ANG5634), XL1-Blue pIMK3-*His-gtcA_K121A_* (ANG5635) and XL1-Blue pIMK3-*His-gtcA_N132A_* (ANG5636). The pIMK3-derivatives were introduced into *L. monocytogenes* strain 10403SΔ*gtcA* by electroporation, resulting in the construction of strains 10403SΔ*gtcA* pIMK3-*gtcA* (ANG5031), 10403SΔ*gtcA* pIMK3-*gtcA_A65S_* (ANG5639), 10403SΔ*gtcA* pIMK3-*gtcA_N69A_* (ANG5640), 10403SΔ*gtcA* pIMK3-*gtcA_V73A_* (ANG5641), 10403SΔ*gtcA* pIMK3-*gtcA_F74A_* (ANG5642), 10403SΔ*gtcA* pIMK3-*gtcA_F91A_* (ANG5643), 10403SΔ*gtcA* pIMK3-*gtcA_R95A_* (ANG5644), 10403SΔ*gtcA* pIMK3-*gtcA_K121A_* (ANG5645), 10403SΔ*gtcA* pIMK3-*gtcA_N132A_* (ANG5646), 10403SΔ*gtcA* pIMK3-*His-gtcA* (ANG5647), 10403SΔ*gtcA* pIMK3-*His-gtcA_A65S_* (ANG5648), 10403SΔ*gtcA* pIMK3-*His-gtcA_N69A_* (ANG5649), 10403SΔ*gtcA* pIMK3-*His-gtcA_V73A_* (ANG5650), 10403SΔ*gtcA* pIMK3-*His-gtcA_F74A_* (ANG5651), 10403SΔ*gtcA* pIMK3-*His-gtcA_F91A_* (ANG5652), 10403SΔ*gtcA* pIMK3-*His-gtcA_R95A_* (ANG5653), 10403SΔ*gtcA* pIMK3-*His-gtcA_K121A_* (ANG5654) and 10403SΔ*gtcA* pIMK3-*His-gtcA_N132A_* (ANG5655).

For the construction of a *B. subtilis gtcA* deletion strain, 1kb-DNA fragments up- and downstream of *gtcA* were amplified using primers ANG3068/3069 and ANG3070/3071, respectively. The resulting PCR products were cut with ApaI and XhoI, respectively and ligated with a Kan cassette, which was excised from pCN34 (ANG201) using ApaI and XhoI. The purified ligation product was transformed into *B. subtilis* 168 (wt), and transformants selected on LB agar plates containing kanamycin. The replacement of *gtcA* with the kanamycin marker was verified by PCR resulting in the construction of *B. subtilis* strains 168Δ*gtcA::kan* (ANG5047). For the complementation of the *B. subtilis gtcA* deletion strain, plasmid pDG1662-P_ywcC_-*ywcC-gtcA* was constructed. To this end, *ywcC*, the first gene of the operon encoding *gtcA*, and *gtcA* were amplified together with the native P_ywcC_ promoter region using primers ANG3089/3090, the PCR product digested with BamHI and HindIII and ligated with plasmid pDG1662, that had been cut with the same enzymes. The resulting plasmid pDG1662-P_ywcC_-*ywcC-gtcA* was recovered in *E. coli* XL1-Blue yielding strain ANG5093. pDG1662-P_ywcC_-*ywcC-gtcA* was linearized using XhoI and transformed into *B. subtilis*Δ*gtcA::kan* yielding the *gtcA* complementation strain *B. subtilis*Δ*gtcA::kan amyE*::P_ywcC_-*ywcC-gtcA* (ANG5102).

### 2.3 GlcNAc staining with WGA

Overnight cultures of *L. monocytogenes* strains were diluted 1:100 in 5 ml BHI medium and grown for 4 h at 37°C until mid-logarithmic growth phase. Bacteria from 100 μl culture were collected by centrifugation for 1 min at 17,000xg. The cell pellet was resuspended in 100 μl PBS pH7.4, mixed with 50 μl of a 0.1 mg/ml Wheat Germ Agglutinin (WGA)-Alexa Fluor^®^ 594 conjugate lectin solution (Invitrogen) and incubated for 5 min at room temperature. The cells were subsequently washed twice with PBS and suspended in 50 μl PBS. 1-1.5 μl of the different samples were spotted on microscope slides that were coated with a thin agarose film (1.2% agarose in distilled water), air-dried and covered with a cover lid. Phase contrast and fluorescence images were taken using a 100x objective and a Zeiss Axio Imager.A1 microscope coupled to the AxioCam MRm and processed using the Zen 2012 (blue edition) software. For the detection of fluorescence signals, the Zeiss filter set 00 was used.

### 2.4 Preparation of cell extract and western blot analysis

For the assessment of LTA production by western blot, *B. subtilis* 168 (wt) and derivatives thereof were grown for 16-20 h in 5 ml LB medium at 30°C. Bacteria from 4 ml culture were collected by centrifugation for 30 min at 17,000xg and bacterial pellets suspended to an OD_600_ of 12 in 100 μl 2x SDS-PAGE sample buffer. The samples were boiled for 45 min, centrifuged for 5 min and 20 μl loaded onto 15% SDS-PAGE gels. *L. monocytogenes* 10403S (wt) and derivatives thereof were grown overnight in 5 ml BHI medium at 37°C. Where indicated, 1 mM IPTG was added to the growth to induce the expression of *gtcA* or *His-gtcA* and the different variants from the complementation plasmid pIMK3. *L. monocytogenes* cell extracts for the detection of LTA were prepared as described previously (Webb et al., 2009). LTA produced by *B. subtilis* and *L. monocytogenes* strains were detected using a polyglycerolphosphate-specific antibody (Clone 55 from Hycult biotechnology) and an HRP-conjugated anti-mouse IgG (Cell Signaling Technologies, USA) at 1:4,000 and 1:10,000 dilutions, respectively. Western blots were developed by the enhanced chemiluminescence method and the signal detected using a ChemiDoc Touch Imager (Bio-Rad). All experiments were performed at least three times and representative images are shown.

For the detection of His-GtcA and its derivatives, bacteria from 20 ml overnight cultures were harvested by centrifugation and OD_600_ readings taken from the same overnight cultures for normalization purposes. The cell pellets were resuspended in 1 ml ZAP buffer containing a protease inhibitor (50 mM Tris-HCl pH 7.5, 200 mM NaCl, 1 complete tablet per 50 ml buffer (Roche)) and cells disrupted three times for 45 sec at 6 m/s using an MP Biomedicals™ Fastprep-24 machine. The cell suspensions were then centrifuged for 15 min at 17,000xg. The resulting pellets were resuspended in 2x SDS-PAGE normalized to an OD_600_ of 40 per 100 μl sample buffer. Samples were incubated for 5 min at 37°C and 25 μl loaded on a 15% Tricine SDS-PAGE gel (Schägger and von Jagow, 1987). For the detection of the His-tagged GtcA proteins, a monoclonal anti-polyHistidine-Peroxidase antibody (Sigma) was used at a 1:10,000 dilution.

### 2.5 LTA and WTA isolation

For the isolation of LTA from *B. subtilis*, the different strains were grown overnight in 2 L LB medium and cells collected by centrifugation. LTA was purified and analyzed using one dimensional (1D) ^1^H nuclear magnetic resonance (NMR) as described previously (Gründling and Schneewind, 2007b, Rismondo et al., 2018). A modified protocol was used for the isolation of LTA from *L. monocytogenes*. Briefly, the strains were grown overnight in 1 L BHI medium that was supplemented with 1 mM IPTG when required. LTA was extracted with butanol and the extracts subsequently dialyzed against water for several days, lyophilized in D_2_O and analyzed by NMR. WTA was purified and analyzed by NMR as previously described (Reichmann et al., 2013, Rismondo et al., 2018).

### 2.6 NMR analysis of cell wall polymers

To analyze the LTA and WTA polymers by ^1^H NMR, 2 mg LTA or 5 mg WTA were suspended and lyophilized twice in 500 μl D_2_O of 99.96% purity. In the final step, LTA and WTA were suspended in 500 μl D_2_O of 99.96% purity and NMR spectra were recorded on a 600-MHz Bruker Advance III spectrometer equipped with a TCl cryoprobe. NMR spectra were recorded at 303 K with a total recycling time of 5 s and a ^1^H flip angle of approximately 30°. Two independent LTA and WTA extractions were performed for each strain. The spectra were annotated according to previously published NMR spectra (Morath et al., 2001, Morath et al., 2002a, Reichmann et al., 2013, Percy et al., 2016, Rismondo et al., 2018). For the calculation of the ratio of GlcNAc to rhamnose modifications on WTA, the area under the peaks at 5.1 ppm and 5 ppm corresponding to one proton in GlcNAc and rhamnose, respectively, were integrated. For the ratio of Galactose:D-alanine modifications on LTA, the area under the peaks at 5.2 ppm and 4.3 ppm corresponding to one proton in galactose and the C-H of D-alanine, respectively, were integrated and the ratio calculated.

### 2.7 Ultra-high performance liquid chromatography tandem mass spectrometry (UPLC-MS/MS)

Purified LTA polymers were depolymerized into monomeric repeating units by hydrolysis of the phosphodiester bonds using 48% hydrofluoric acid for 20 h at 0°C. The LTA monomers were then lyophilized and subjected to UPLC-MS/MS analysis as previously described (Shen et al., 2017). All data were collected and processed using the MassLynx software, version 4.1 (Waters Corp., USA), and MS spectra were background-corrected by subtracting the signals between 0–1 min of their respective chromatograms.

## 3. RESULTS

### 3.1 The predicted *B. subtilis* glycosyltransferases YfhO is sugar- and acceptor-specific

The *B. subtilis* enzymes CsbB and YfhO have recently been shown to be required for the decoration of LTA with GlcNAc residues (Fig. 1C) (Rismondo et al., 2018). YfhO likely acts as extracellular GT mediating the transfer of GlcNAc from the C_55_-P-GlcNAc lipid intermediate on to the LTA backbone (Rismondo et al., 2018). In contrast Lmo1079, the YfhO homolog of *L. monocytogenes*, is necessary for the modification of WTA with GlcNAc residues (Fig. 1A) (Rismondo et al., 2018, Denes et al., 2015, Eugster et al., 2015). This suggests that *B. subtilis* and *L. monocytogenes* YfhO enzymes use the same C_55_-P-GlcNAc sugar molecule as substrate, however, they use LTA and WTA, respectively, as acceptor molecules. To further test the sugar and acceptor molecule specificity of the *B. subtilis* YfhO enzymes, the *B. subtilis csbB-yfhO* operon was expressed in *L. monocytogenes* strains lacking sugar modifications either on LTA (strain 10403SΔ*gtlAB*) or sugar modifications on WTA (strain 10403SΔ*lmo1079*Δ*lmo2550*). Subsequently, the structures of WTA and LTA were analyzed by NMR and western blot. As expected, the WTA polymer isolated from the wildtype *L. monocytogenes* strain 10403S was decorated with GlcNAc residues and this modification was absent in strain 10403SΔ*lmo1079*Δ*lmo2550* (Fig. S1). The WTA polymer produced by *L. monocytogenes* strain 10403SΔ*lmo1079* Δ*lmo2550*+*csbB-yfhO* was indistinguishable from the polymer produced by the *lmo1079/lmo2550* mutant, revealing that the *B. subtilis* CsbB and YfhO enzymes are unable to glycosylate the WTA of *L. monocytogenes*. Next, cell extracts of *L. monocytogenes* strains 10403S, 10403SΔ*gtlAB* and 10403SΔ*gtlAB*+*csbB-yfhO* were analyzed by western blot using a polyglycerolphosphate specific antibody, which recognizes the LTA backbone. In previous studies it has been shown that the absence of sugar modifications on LTA results in a stronger LTA signal on western blots (Rismondo et al., 2018, Percy et al., 2016). Consistent with these findings, the signal was increased for the extracts isolated from the *gtlAB* mutant as compared to the wildtype strain (Fig. 2A). In contrast, extracts isolated from strain 10403SΔ*gtlAB*+*csbB-yfhO* produced a low signal (Fig. 2A), suggesting that the LTA of strain 10403SΔ*gtlAB*+*csbB-yfhO* is again modified with sugar residues. To investigate this further, LTA was isolated from wildtype 10403S, the *gtlAB* mutant and strain 10403SΔ*gtlAB*+*csbB-yfhO* and analyzed by 1D ^1^H NMR and mass spectrometry. Galactose-specific peaks (colored yellow in Fig. 2) could be detected for the LTA isolated from the wildtype strain 10403S and were absent in the NMR spectra obtained for the LTA of the *gtlAB* mutant and *gtlAB* mutant expressing *csbB-yfhO* (Fig. 2B-C). In contrast, four additional peaks could be detected in the NMR spectra obtained for the LTA isolated from strain 10403SΔ*gtlAB*+*csbB-yfhO* as compared to the LTA derived from the *gtlAB* mutant (Fig. 2D). The chemical shifts of these additional peaks differ from those obtained for the galactose residues and resembled those observed for GlcNAc residues present on *B. subtilis* LTA (Rismondo et al., 2018). To verify that the LTA produced by strain 10403SΔ*gtlAB*+*csbB-yfhO* is indeed decorated with GlcNAc residues, LTA isolated from strains 10403S, 10403SΔ*gtlAB* and 10403SΔ*gtlAB*+*csbB-yfhO* was depolymerized with hydrofluoric acid and analyzed by UPLC-MS. For the wildtype *L. monocytogenes* strain 10403S, a peak with an m/z of 253.09 was detected corresponding to galactose-glycerol moieties. In contrast, a peak with an m/z of 294.12 was observed for the depolymerized LTA produced by strain 10403SΔ*gtlAB*+*csbB-yfhO*, consistent with GlcNAc-glycerol moieties. As expected, neither of these two peaks was detected for the LTA sample isolated from the *gtlAB* mutant (Fig. S2). Taken together, these results show that the putative glycosyltransferases CsbB and YfhO specifically glycosylate LTA with GlcNAc residues regardless if they are expressed in *L. monocytogenes* or *B. subtilis*.

**Figure 2:**
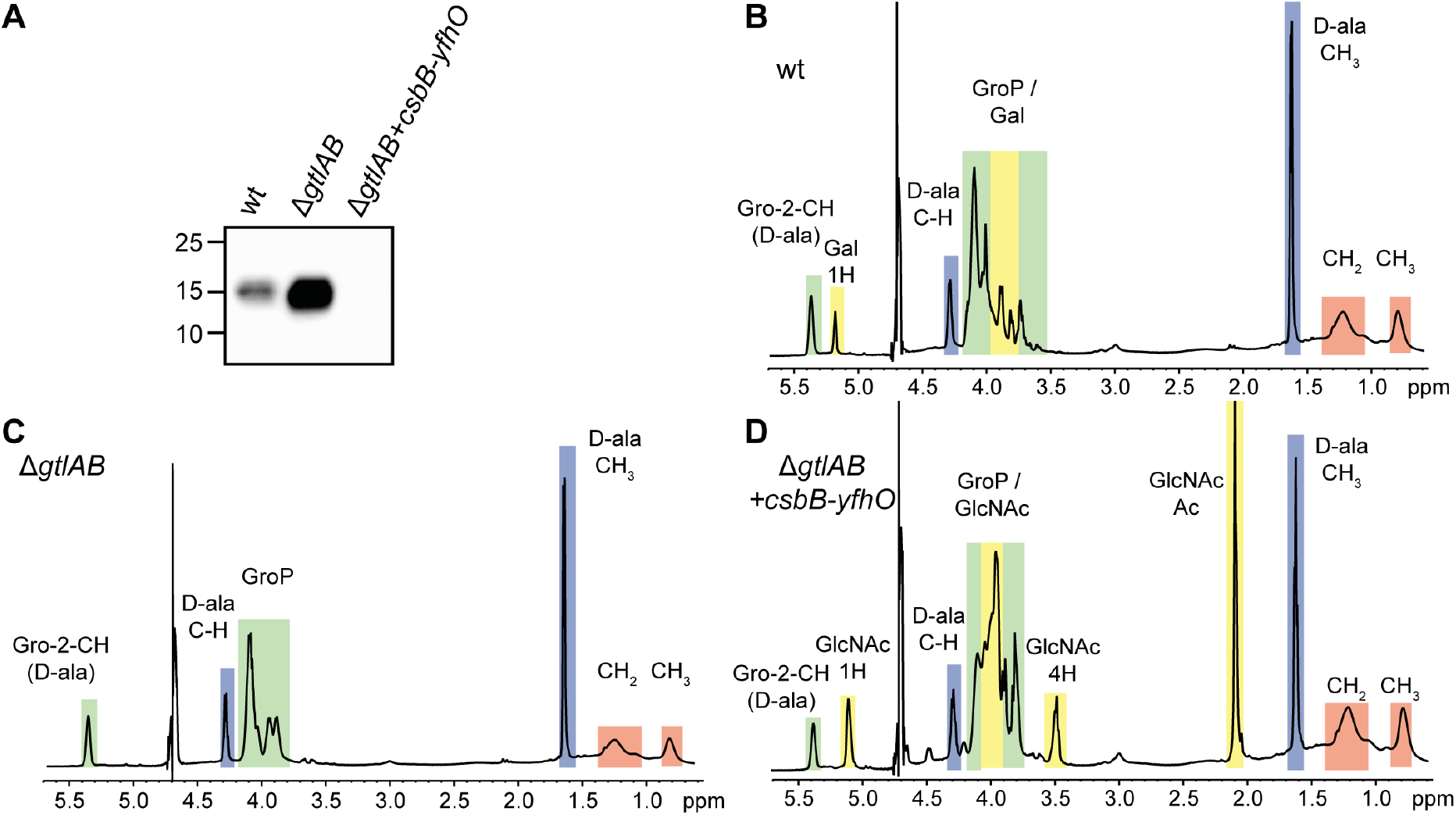
LTA in *L. monocytogenes* 10403S is glycosylated with GlcNAc residues upon expression of *B. subtilis* CsbB and YfhO. (A) Detection of LTA by western blot. Cell extracts of *L. monocytogenes* strains 10403S (wt), 10403SΔ*gtlAB* and 10403SΔ*gtlAB*+*csbB-yfhO* were prepared and separated on a 15% SDS-PAGE gel. LTA was detected by western blot using a polyglycerolphosphate-specific monoclonal antibody. (B-D) NMR spectra of LTA isolated from *L. monocytogenes* strains 10403S (B), 10403SΔ*gtlAB* or (C) 10403SΔ*gtlAB*+*csbB-yfhO* (D). Colored boxes and labels indicate nonexchangeable protons derived from the different LTA components. Peaks were assigned as previously described (Wörmann et al., 2011, Morath et al., 2001, Morath et al., 2002a, Morath et al., 2002b). The spectra are representatives of three independent experiments.

### 3.2 GtcA is required for the glycosylation of LTA in *L. monocytogenes* and *B. subtilis*

GtlA and CsbB have been identified as putative cytoplasmic GTs and GtlB and YfhO as GTs with extracellular activity involved in the glycosylation of LTA in *L. monocytogenes* 10403S and *B. subtilis* 168, respectively (Fig. 1) (Percy et al., 2016, Rismondo et al., 2018). However, the enzyme involved in the flipping of the lipid-linked sugar intermediate has not been identified in these two organisms. In Gram-negative bacteria, members of the Wzx family have been identified as flippases of C_55_-P-linked oligosaccharides such as the C_55_-P-linked O-antigen subunits (Islam and Lam, 2013, Islam and Lam, 2014). In *Streptococcus pneumoniae*, Wzx is involved in the transport of the final subunit produced during capsule synthesis (Robbins et al., 1966, Xayarath and Yother, 2007). Using the *S. pneumoniae* Wzx sequence (locus tag AF316641_9) as a query in a BLASTP search, Lmo0215 was identified as the closest Wzx homolog in *L. monocytogenes* with an amino acid identity of 26%. In addition to Wzx transporters, members of the GtrA protein family such as GtrA of *Shigella flexneri* and Rv3789 of *Mycobacterium tuberculosis* are thought to be involved in the flipping of lipid-linked sugar intermediates (Korres et al., 2005, Larrouy-Maumus et al., 2012). In a previous study, it was shown that GtcA (Lmo2549), a GtrA protein family member, is involved in the glycosylation of WTA in *L. monocytogenes* (Promadej et al., 1999). To determine if either GtcA or the Wzx homolog Lmo0215 is required for the glycosylation of LTA in *L. monocytogenes* 10403S, *gtcA* and *lmo0215* mutants were constructed. Next, cell extracts were prepared from wildtype 10403S, the *gtcA* and *lmo0215* deletion strains and analyzed by western blot using a polyglycerolphosphate-specific LTA antibody. No difference in signal intensities was observed between the extracts isolated from the wildtype and Δ*lmo0215* mutant strains, suggesting that the encoded protein is not involved in the LTA glycosylation process. In contrast, an increased LTA signal was observed for the *gtcA* mutant strain (Fig. 3A). The signal was of similar intensity to that observed for extracts derived from *gtlA* and *gtlB* mutants, strains known to lack galactose modifications on their LTA (Fig. 3B). This phenotype could be complemented by expressing *gtcA* from an IPTG-inducible promoter (Fig. 3B). Indeed, partial complementation and a reduction in LTA signal could already be seen in the absence of inducer, indicating basal-level expression of *gtcA* even in the absence of IPTG (Fig. 3A). To investigate the involvement of GtcA in the LTA glycosylation process further, LTA was isolated from wildtype 10403S, the *gtcA* deletion and complementation strain (grown in the presence of 1 mM IPTG) and analyzed by 1D ^1^H NMR. This analysis showed that LTA of the Δ*gtcA* deletion strain lacks the galactose-specific peaks (Fig. 4B), whereas the LTA derived from strains 10403S and the *gtcA* complementation strain were glycosylated (Fig. 4A+C).

**Figure 3:**
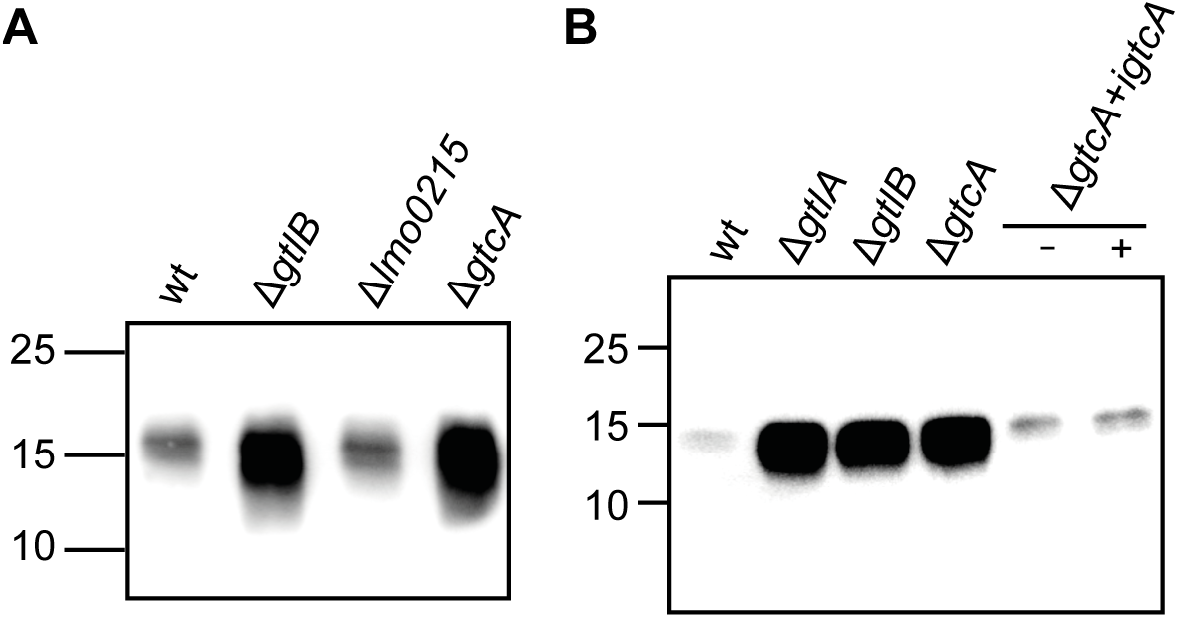
LTA production in *L. monocytogenes* wildtype, *gtcA* and *lmo0215* mutant strains. (A-B) Detection of LTA by western blot. Cell extracts of *L. monocytogenes* strains (A) 10403S (wt), the *gtcA* and *lmo0215* mutants or (B) 10403S (wt), the *gtcA* mutant and the *gtcA* complementation strain grown in the absence (-) or presence (+) of IPTG were prepared and separated on a 15% SDS-PAGE gel. LTA was detected by western blot using a polyglycerol-phosphate-specific monoclonal antibody. *L. monocytogenes gtlA* and *gtlB* mutant strains, defective in LTA glycosylation (Percy et al., 2016, Rismondo et al., 2018) were included as positive controls. One representative result of three independent experiments is shown.

**Figure 4:**
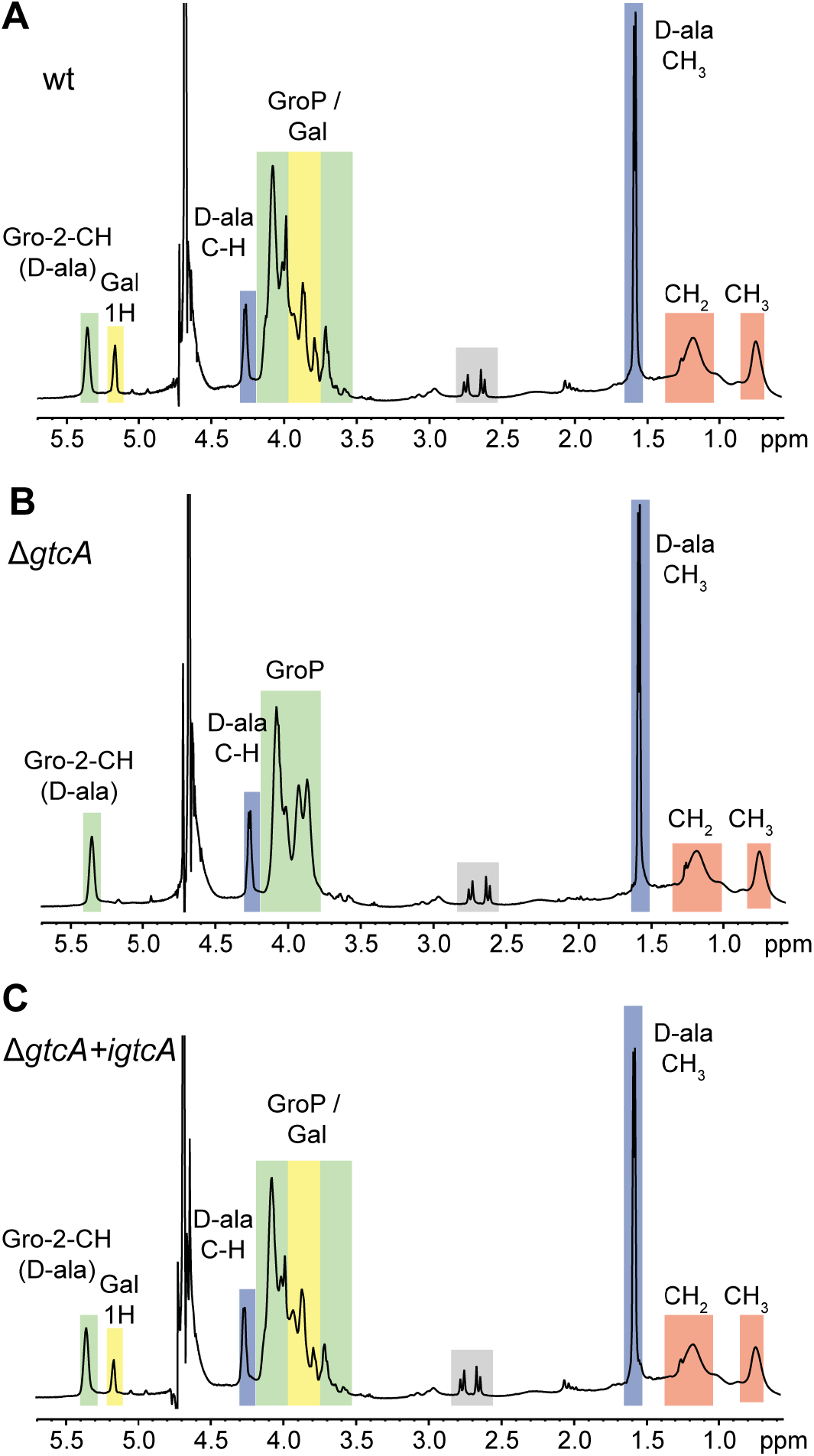
NMR analysis of LTA isolated from wildtype *L. monocytogenes* strain 10403S, and isogenic *gtcA* mutant and complementation strains. (A-C) NMR spectra of LTA produced by *L. monocytogenes* strains (A) 10403S (wt), (B) the *gtcA* mutant and (C) the *gtcA* complementation strain (grown in presence of IPTG). Peaks of nonexchangeable protons were assigned to the different LTA components according to previously published spectra and are highlighted in colored boxes (Morath et al., 2001, Morath et al., 2002a, Morath et al., 2002b, Wörmann et al., 2011). The spectra are representatives of two independent experiments.

Taken together, these data highlight that GtcA is not only required for the WTA glycosylation process in *L. monocytogenes* 10403S but also needed for the glycosylation of LTA.

In *B. subtilis* 168, *gtcA* is part of the *ywcC-gtcA-galK-galT* operon, also encoding the TetR-like transcriptional regulator YwcC, the galactokinase GalK and the galactose-1-phosphate uridylyltransferase GalT (Fig. 5A). To test if GtcA is also involved in the LTA glycosylation process, *gtcA* deletion and complementation strains were constructed and LTA production assessed by western blot. A stronger signal was observed for extracts derived from the *gtcA* mutant compared to the wildtype 168 strain, which was similar to that of the *csbB* mutant control strain, which is known to lack GlcNAc modification on LTA (Rismondo et al., 2018) (Fig. 5B). The LTA signal was reduced back to wildtype levels in the *gtcA* complementation strain 168Δ*gtcA*+*ywcC-gtcA*, in which *gtcA* was expressed along with *ywcC*, the upstream gene, from its native promoter (Fig. 5B). These data indicate that *gtcA* is also required for the LTA glycosylation process in *B. subtilis* 168. This was further confirmed by NMR analysis of LTA isolated from the wildtype, *gtcA* mutant and *gtcA* complementation strains, where GlcNAc-specific peaks could be detected for the wildtype and complementation strain but not for the *gtcA* mutant (Fig. 5C-E). Taken together, our results show that GtcA is involved in the LTA glycosylation in both, *B. subtilis* 168 and *L. monocytogenes* 10403S, and potentially acts as a flippase to transport the C_55_-P-sugar intermediate across the membrane.

**Figure 5:**
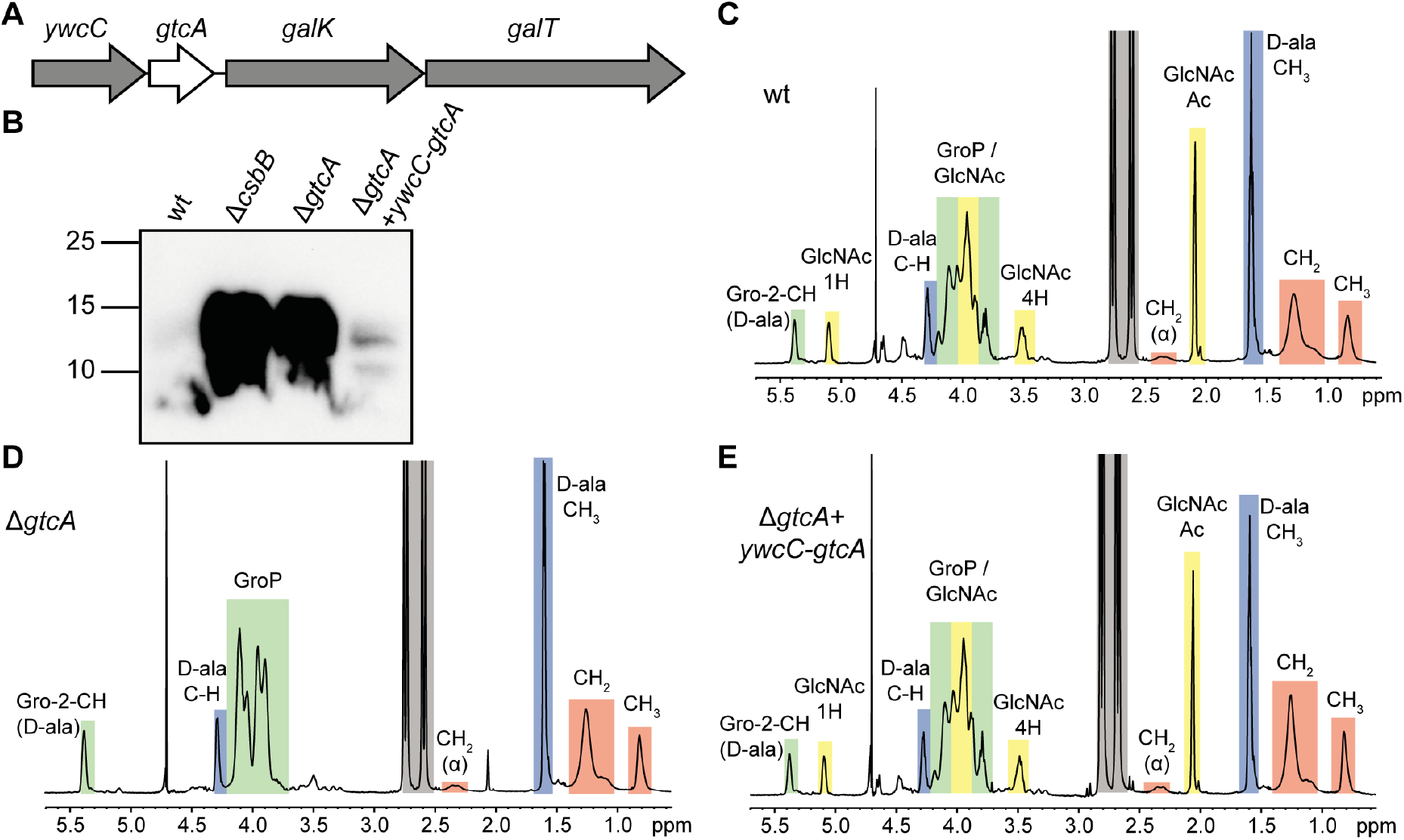
LTA production and NMR analysis of LTA isolated from wildtype *B. subtilis* 168 and the isogenic *gtcA* mutant and complementation strains. (A) Schematic representation of the *ywcC-gtcA-galK-galT* operon. (B) Analysis of LTA by western blot. Cell extracts of *B. subtilis* strain 168 (wt), the *gtcA* mutant and the *gtcA* complementation strain (*gtcA*+*ywcC-gtcA*) were prepared and separated on a 15% SDS-PAGE gel. LTA was detected by western blot using a polyglycerol-phosphate-specific monoclonal antibody. Cell extract of strain *csbB* was included as positive control (Rismondo et al., 2018). One representative result of four independent experiments is shown. (C-E) NMR spectra of LTA produced by (C) *B. subtilis* 168 (wt), (D) the *gtcA* mutant and (E) the *gtcA* complementation strain. Peaks of nonexchangeable protons were assigned to the different LTA components based on previously published spectra and highlighted in colored boxes (Wörmann et al., 2011, Morath et al., 2001, Morath et al., 2002a, Morath et al., 2002b). The spectra are representatives of two independent experiments.

### 3.3 Identification of amino acids essential for the function of GtcA

GtrA family proteins are thought to act as flippases of lipid-linked sugar intermediates in different bacteria (Larrouy-Maumus et al., 2012, Kho and Meredith, 2018, Korres et al., 2005, Promadej et al., 1999). However, so far it is not known which amino acids play an important role for the function of these proteins. Based on the membrane topology prediction using the TMHMM server 2.0 (Sonnhammer et al., 1998), the *L. monocytogenes* GtcA protein contains four transmembrane helices (Fig. 6A). To identify conserved amino acid residues, the *L. monocytogenes* GtcA protein sequence was used in a BLASTP search against the non-redundant protein data base, the sequences of the top 1000 homologs were aligned using Jalview (Waterhouse et al., 2009) and a WebLogo generated (Crooks et al., 2004) (Fig. 6B). Eight amino acids, A65, N69, V73, F74, F91, R95, K121 and N132 located in different parts of the *L. monocytogenes* GtcA protein (Fig. 6A), were found to be conserved in 99-100% of the analyzed proteins. To determine, if any of these highly conserved amino acid residues are important for the activity of GtcA, GtcA and derivatives harbouring single amino acid substitutions were expressed from an IPTG-inducible promoter in the *L. monocytogenes gtcA* deletion strain. The LTA glycosylation state of strains expressing the different GtcA variants was assessed by western blot (Fig. 7A) and the WTA glycosylation state was assessed by fluorescence microscopy using the fluorescently labelled lectin wheat germ agglutinin (WGA)-Alexa Fluor^®^ 594 conjugate, which recognizes GlcNAc residues on WTA (Loessner et al., 2002) (Fig. 7B). Wildtype 10403S and the *gtcA* mutant strains were used as controls in these experiments (Fig. 7). Based on this analysis, the GtcA variants with mutations in the highly conserved residues could be grouped into four different categories: GtcA variants with unaltered function (A65S, N69A, V73A), GtcA variants, which showed a defect in both LTA and WTA glycosylation (R95A, N132A), a GtcA variant which was defective in LTA glycosylation (K121A) and GtcA variants with a defect in WTA glycosylation (F74A, F91A) (Fig. 7, Table S3). To determine whether the GtcA derivatives GtcA_F74A_, GtcA_F91A_, GtcA_R95A_, GtcA_K121A_ and GtcA_N132A_ fail to participate in the glycosylation processes of LTA and/or WTA due to differences in protein expression, GtcA and derivatives thereof where expressed in *L. monocytogenes* strain 10403SΔ*gtcA* as His-tag fusion proteins. *L. monocytogenes* strain 10403SΔ*gtcA*+*His-gtcA* produced glycosylated LTA and WTA, suggesting that the N-terminal His-tag does not ablate the function of the GtcA protein (Fig. S3 and S4). Similar results were obtained for the *gtcA* deletion strain expressing His-GtcA variants carrying point mutations in the conserved amino acids as compared to the untagged GtcA variants when the LTA and WTA glycosylation status was assessed by western blot and fluorescence microscopy, respectively (Fig. S3A+C, Table S3). Next, protein extracts were prepared from control and *L. monocytogenes* strains expressing the different His-GtcA variants and the His-tagged proteins detected by western-blot. All His-GtcA proteins could be detected and most proteins were produced at similar amounts as the wildtype His-GtcA control protein, with exception of His-GtcA_F74A_ and His-GtcA_R95A_, which were produced at slightly lower levels (Fig. S3B). Since the GtcA variants with the F74A or R95A mutations also showed a defect in function (Fig. S3A+C, Table S3), we wanted to determine whether the observed reduction in protein production could be the reason for this as opposed to the actual amino acid substitution. To assess the impact of reduced GtcA production on LTA and WTA glycosylation, *L. monocytogenes* strain 10403SΔ*gtcA*+*His-gtcA* was grown in the absence of IPTG and LTA and WTA glycosylation assessed. This analysis revealed that basal expression of His-GtcA is sufficient to fully reduce the LTA western-blot signal to that of a wild-type strain and partially restore the glycosylation of WTA as assessed by fluorescence microscopy, even though no protein could be detected by western blot (Fig. S4). These results indicate that amino acid residues F74 and R95 likely play an important role for the function of GtcA as the reduced protein production alone cannot explain the observed phenotypes.

**Figure 6:**
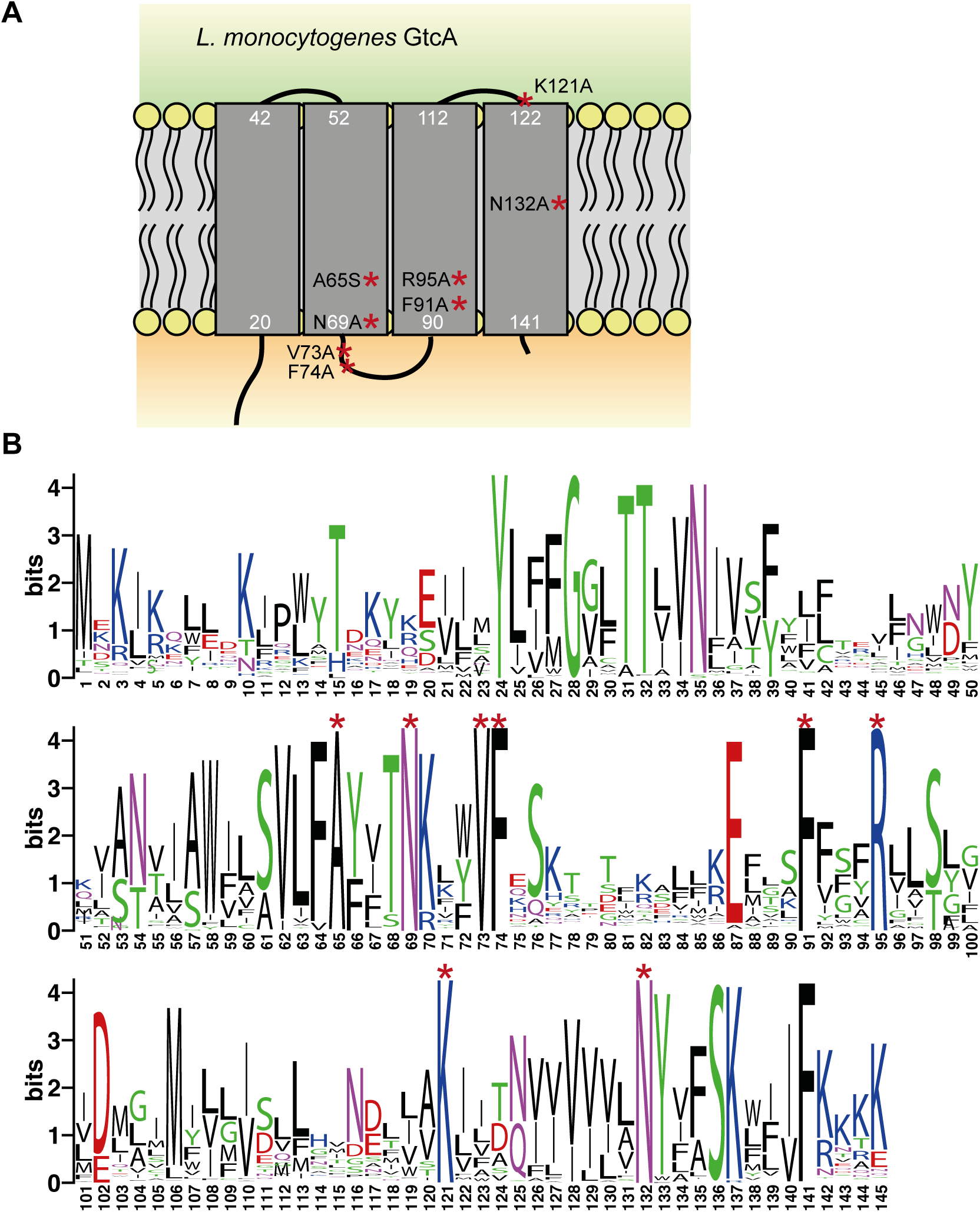
Membrane topology model and amino acid conservation of the *L. monocytogenes* 10403S GtcA protein. (A) Membrane topology model of the *L. monocytogenes* protein GtcA based on the prediction using the TMHMM version 2 server (Sonnhammer et al., 1998). Conserved amino acids mutated as part of this study are indicated. The numbers shown in white indicate the amino acids located at the predicted boarders of the TM helices. (B) WebLogo motif of GtcA proteins. *L. monocytogenes* GtcA was used as query sequence in a BLASTP search and the sequences of the top 1000 GtcA homologs downloaded from the NCBI website (on 21.03.2019) and aligned using Jalview (Waterhouse et al., 2009). The alignment was used to generate the presented WebLogo motif (Crooks et al., 2004).

**Figure 7:**
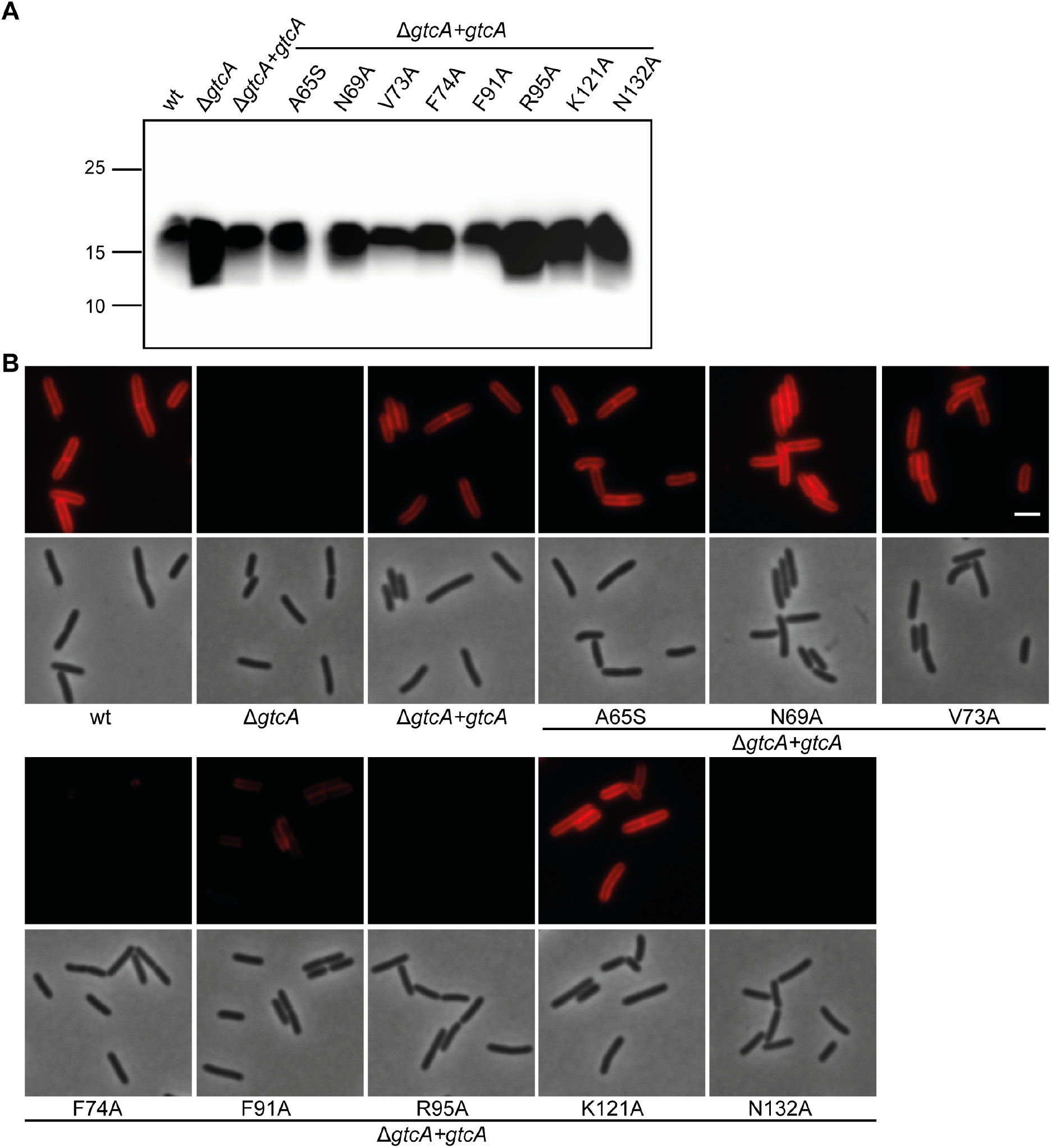
Identification of amino acid residues essential for the function and/or stability of the *L. monocytogenes* 10403S GtcA protein. (A) Detection of LTA by western blot. Cell extracts of the indicated *L. monocytogenes* strains were prepared, separated on a 15% SDS PAGE and LTA detected using a polyglycerolphosphate-specific monoclonal antibody. (B) Microscopy analysis and detection of WTA glycosylation using the fluorescently labelled WGA-Alexa 594 lectin. Log-phase cells of the indicated strains were stained with WGA-Alexa 594 as described in the methods section and subjected to phase and fluorescence microscopy. Scale bar is 2 μm. One representative result of three independent experiments is shown.

Through the bioinformatics, mutagenesis, western blot and fluorescent microscopy analysis, we identified key amino acids in GtcA that appear to play an important role for the glycosylation of LTA but not WTA glycosylation (K121) and vice versa (F74 and F91). To determine the actual chemical structure of WTA and LTA produced by such variants, LTA and WTA were isolated from strains 10403SΔ*gtcA*+*gtcA*, 10403SΔ*gtcA*+*gtcA_F74A_* and 10403SΔ*gtcA*+*gtcA_K121A_*, and analyzed by 1D ^1^H NMR. The NMR spectra of WTA extracted from strains 10403SΔ*gtcA*+*gtcA* and 10403SΔ*gtcA*+*gtcA_K121A_* were comparable to each other (Fig. 8A+C). In contrast, the GlcNAc-specific peaks were significantly smaller in the NMR spectra for the WTA produced by strain 10403SΔ*gtcA*+*gtcA_F74A_* (Fig. 8B). To quantify the structural differences, the ratio of GlcNAc and rhamnose modifications on WTA of the different strains was calculated as described in the methods section. WTA of *L. monocytogenes* strains 10403SΔ*gtcA*+*gtcA* and 10403SΔ*gtcA*+*gtcA_K121A_* have a GlcNAc:rhamnose ratio of around 0.7, whereas a GlcNAc:rhamnose ratio of 0.16 was observed for WTA for strain 10403SΔ*gtcA*+*gtcA_F74A_* (Fig. 8D). Galactose residues could be detected on the LTA produced in all three *L. monocytogenes* strains (10403SΔ*gtcA*+*gtcA*, 10403SΔ*gtcA*+*gtcA_F74A_* and 10403SΔ*gtcA*+*gtcA_K121A_*) (Fig. 8E-G). However, the calculation of the galactose:D-alanine ratio (1H galactose peak at 5.2 ppm : C-H D-alanine peak at 4.3 ppm) indicated a slight increase in glycosylation of LTA in strain 10403SΔ*gtcA*+*gtcA_F74A_* and as expected a decrease in LTA glycosylation in strain 10403SΔ*gtcA*+*gtcA_K121A_* as compared to the LTA isolated from strain 10403SΔ*gtcA*+*gtcA* (Fig. 8H). Taken together, these data are consistent with the changes observed in the western blot and microscopy analysis. They indicate a predominant requirement of residue F74 in GtcA for the glycosylation of WTA with GlcNAc residues without leading to a reduction in the glycosylation of LTA with galactose. In contrast, residue K121 is important for the glycosylation of LTA but is not essential for WTA glycosylation.

**Figure 8:**
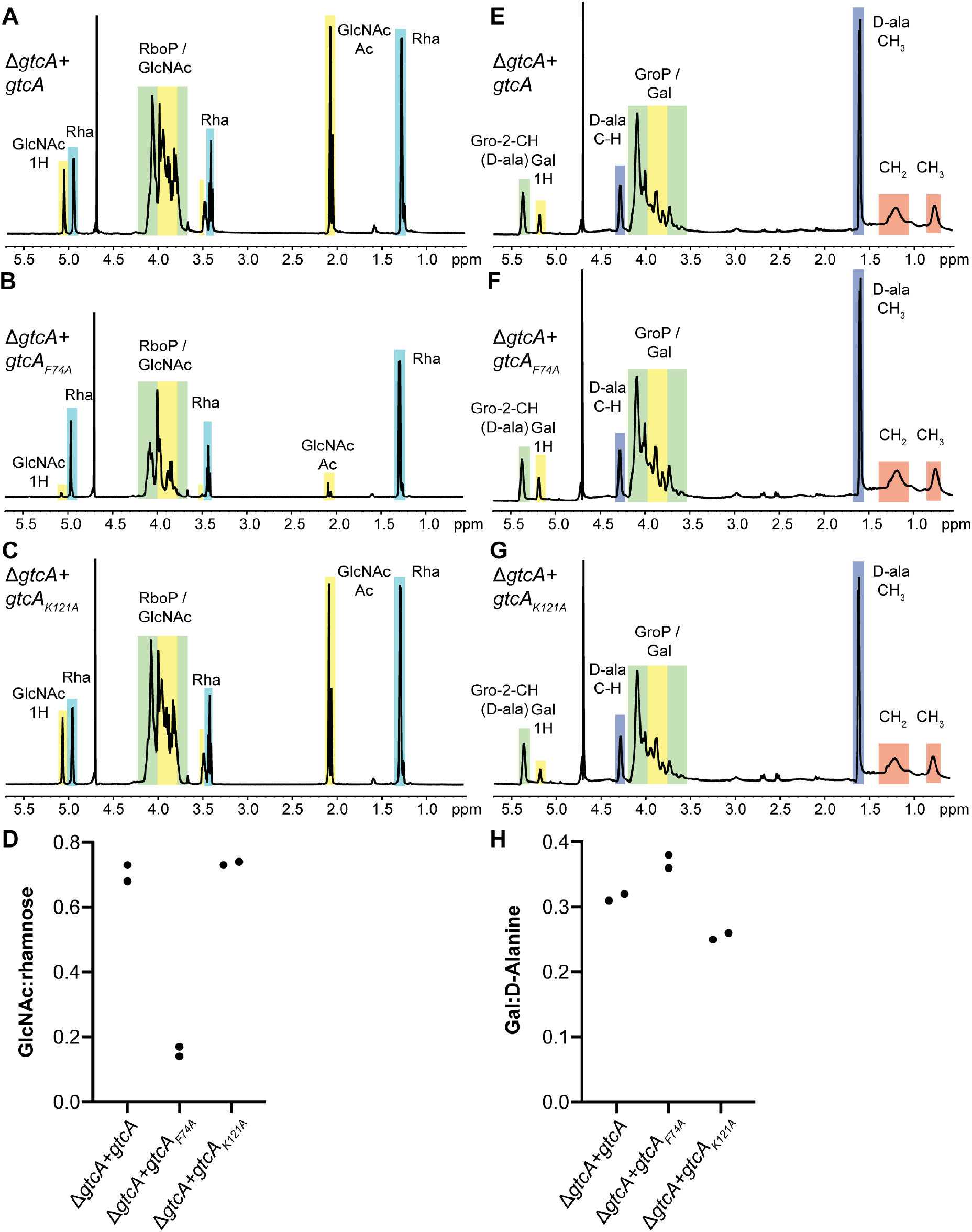
Quantification of LTA and WTA sugar modifications in *L. monocytogenes* 10403S producing wildtype GtcA or the GtcA_F74A_ or GtcA_K121A_ variants. (A-C) NMR spectra of WTA. WTA was extracted from *L. monocytogenes* strains (A) 10403SΔ*gtcA*+*gtcA*, (B) 10403SΔ*gtcA*+*gtcA_F74A_* and (C) 10403SΔ*gtcA*+*gtcA_K121A_* and analyzed by NMR. The spectra are representatives of two independent experiments. (D) Ratio of GlcNAc and rhamnose modifications on WTA. The peaks at 5.1 ppm and 5 ppm corresponding to 1H of GlcNAc and rhamnose, respectively, were integrated and the ratio of GlcNAc:rhamnose calculated and plotted for the two independent WTA extractions. (E-G) NMR spectra of LTA. LTA was extracted from *L. monocytogenes* strains (E) 10403SΔ*gtcA*+*gtcA*, (F) 10403SΔ*gtcA*+*gtcA_F74A_* and (G) 10403SΔ*gtcA*+*gtcA_K121A_*. The spectra are representatives of two independent experiments. (H) Ratio of Gal and D-alanine modifications on LTA. The peaks at 5.2 ppm and 4.3 ppm corresponding to 1H of galactose and C-H of D-alanine, respectively, were integrated and the ratio of Gal:D-Ala calculated and plotted for the two independent LTA extractions.

## DISCUSSION

Recent studies have provided insight into the protein components forming part of the proposed three-component glycosylation system required for the glycosylation of LTA in *B. subtilis* and *S. aureus* and for the glycosylation of both polymers, LTA and WTA, in the *L. monocytogenes* serovar 1/2a strain 10403S (Kho and Meredith, 2018, Rismondo et al., 2018, Percy et al., 2016). YfhO, a putative glycosyltransferase, is thought to transfer GlcNAc residues from a C_55_-P-GlcNAc intermediate onto the LTA backbone in *B. subtilis* and *S. aureus* (Kho and Meredith, 2018, Rismondo et al., 2018). Interestingly, the *L. monocytogenes* serovar 1/2a YfhO homolog Lmo1079, is required for the decoration of WTA with GlcNAc residues rather than the modification of LTA with galactose residues (Eugster et al., 2015, Rismondo et al., 2018). In the case of *L. monocytogenes* strain 10403S, the predicted glycosyltransferase GtlB is required for the modification of LTA with galactose residues. (Rismondo et al., 2018). While YfhO-like proteins have 12 predicted transmembrane helices and a large outside loop between the last two helices, GtlB-like proteins possess only eight transmembrane helices and a smaller extracellular loop located between the first two helices (Rismondo et al., 2018). Such large sequence differences and the lack of structural information make it hard to bioinformatically predict which glycosyltransferases is required for decoration of teichoic acids and if they use LTA or WTA as acceptor molecules. The observation that YfhO of *B. subtilis* and *S. aureus* and the *L. monocytogenes* 10403S YfhO homolog Lmo1079 are required for the glycosylation of LTA and WTA, respectively (Eugster et al., 2015, Kho and Meredith, 2018, Rismondo et al., 2018), suggests that these enzymes specifically recognize their cognate cell wall polymer. Here, we tested whether the *B. subtilis* enzymes YfhO acts sugar- and acceptor-specific by expressing the *B. subtilis csbB-yfhO* operon in *L. monocytogenes gtlAB* or *lmo2550/lmo1079* deletion strains, which are unable to glycosylate LTA or WTA, respectively. This analysis showed that expression of the *csbB-yfhO* operon leads to the attachment of GlcNAc residues onto the LTA polymer in *L. monocytogenes* suggesting that the *B. subtilis* YfhO enzyme specifically recognizes and transfers GlcNAc residues to the backbone of LTA. Both, *B. subtilis* 168 and *L. monocytogenes* 10403S possess a type I LTA with a GroP backbone (Percy and Gründling, 2014) and our data indicate that the *B. subtilis* YfhO protein recognizes the GroP backbone without the need of any additional specificity factors since we only introduced the *B. subtilis csbB-yfhO* operon into *L. monocytogenes*. It is however unclear if in our experiments the C_55_-P-GlcNAc intermediate used by YfhO is solely produced by the *B. subtilis* CsbB protein or whether YfhO also utilized the C_55_-P-GlcNAc intermediate produced by the *L. monocytogenes* GT Lmo2550, which seems more likely. As discussed in more detail below, the C_55_-P-GlcNAc is likely transported across the membrane by the *L. monocytogenes* GtcA protein.

For the glycosylation of LTA and WTA in *L. monocytogenes*, a C_55_-P-sugar intermediate needs to be transported across the membrane. Several enzymes have been proposed for the translocation of lipid-linked sugar intermediates for different glycosylation processes and biosynthetic pathways. MurJ and Amj have been identified as the enzyme responsible for the flipping of lipid II during peptidoglycan biosynthesis (Fay and Dworkin, 2009, Meeske et al., 2015, Ruiz, 2008, Sham et al., 2014, Inoue et al., 2008). Another class are Wzx (or RfbX) flippases, which are involved in the flipping of lipid carrier linked O-antigen repeating units during LPS biosynthesis in *Escherichia coli*, *Salmonella enterica* and *Shigella flexneri* (Liu et al., 1996, Feldman et al., 1999) or the flipping of lipid carrier linked oligosaccharides during capsule biosynthesis in *Streptococcus pneumoniae* (Bentley et al., 2006). Both, MurJ and Wzx flippases, belong to the multidrug/oligosaccharidyl lipid/polysaccharide (MOP) transporter family of proteins (Hvorup et al., 2003) and have 12-14 transmembrane helices. A third class of proteins involved in the translocation of lipid-linked intermediates are small multidrug resistance (SMR) transporters, which are small hydrophobic proteins with four predicted transmembrane helices (Paulsen et al., 1996). The *E. coli* ArnE/ArnF proteins are members of this family and they are thought to transport a C_55_-P-sugar intermediate across the membrane, which is necessary for the glycosylation of Lipid A (Yan et al., 2007). Another member of the SMR family is the *S. flexneri* SfX bacteriophage protein GtrA (Guan et al., 1999). Indeed, bacteriophage SfX encodes a three-protein glycosylation system comprised of GtrA, GtrB and GtrX (also referred to as Gtr_type_) and these proteins are responsible for serotype conversion in *S. flexneri* (Huan et al., 1997a, Huan et al., 1997b, Mavris et al., 1997). GtrB, a homolog of the *L. monocytogenes* Lmo2550 and GtlA proteins, is necessary for the formation of a C_55_-P-sugar intermediate, which is flipped across the membrane by the putative flippase GtrA and subsequently attached to the O-antigen by GtrX (Guan et al., 1999, Korres et al., 2005, Allison and Verma, 2000). The flippases required for the translocation of the C_55_-P sugar intermediates during LTA glycosylation in *L. monocytogenes* and *B. subtilis* have not been identified thus far. As part of this study, we show that Lmo0215, a *L. monocytogenes* protein showing homology to Wzx-type transporters, is not required for the glycosylation of LTA in *L. monocytogenes*. In contrast, absence of the GtrA homolog GtcA results in the loss of sugar modifications on LTA in *L. monocytogenes* 10403S as well as in *B. subtilis* 168. The same protein has been previously shown to be important for WTA glycosylation in *L. monocytogenes* and *L. innocua* and has been proposed to act as a flippase enzyme (Promadej et al., 1999, Lan et al., 2000). In addition, a recent study suggested that a GtcA homologue is responsible for the translocation of C_55_-P-GlcNAc during the LTA glycosylation process in *S. aureus* (Kho and Meredith, 2018). Members of the GtrA protein family are usually small and highly hydrophobic proteins, which are predicted to contain three to four transmembrane helices (TMs) and their function has been associated with the synthesis of diverse cell surface polysaccharide. In *Mycobacterium smegmatis* and *M. tuberculosis*, the GtrA homolog Rv3789 is involved in the arabinosylation of arabinogalactan and lipoarabinomannan (Larrouy-Maumus et al., 2012, Kolly et al., 2015). However, there is conflicting evidence in the literature concerning the role of Rv3789 during the arabinosylation process. It has been proposed that Rv3789 acts as a flippase enzyme to translocate decaprenyl-phospho-arabinose (DPA) across the cytoplasmic membrane (Larrouy-Maumus et al., 2012). However, it has also been suggested that Rv3789 acts as an anchor protein by recruiting other proteins involved in the arabinogalactan biosynthesis, such as the priming arabinosyltransferase AftA (Kolly et al., 2015, Brecik et al., 2015). Our data show that GtcA is involved in the LTA glycosylation process in *L. monocytogenes* and *B. subtilis* and we favor a model where the protein acts as C_55_-P-sugar flippase enzyme rather than an anchor protein. However actual biochemical evidence for such an activity is still lacking and will need to be addressed in future studies.

Deletion of *gtcA* in *B. subtilis* 168 resulted in a loss of GlcNAc modifications on LTA. It is interesting to note that the *B. subtilis* GtcA protein is encoded in an operon with other genes coding for galactose metabolism proteins (Glaser et al., 1993). The genetic location of *gtcA* might therefore indicate that GtcA is also required for the glycosylation of a different polymer or proteins with galactose or a galactose derivative. Similarly, if the *L. monocytogenes* GtcA protein indeed acts as a flippase enzyme, our results indicate that it has a broader substrate specificity as it impacts glycosylation of LTA and WTA with galactose and GlcNAc residues, respectively (Fig. 4, 7B). Whereas *L. monocytogenes* and *S. aureus* only possess one GtcA protein, two GtrA-like proteins, GtcA and YngA, are encoded in the *B. subtilis* 168 genome. As deletion of *gtcA* alone leads to a complete absence of GlcNAc residues on LTA in *B. subtilis*, this suggests that under the conditions tested YngA is not required for this process. We speculate that *B. subtilis* YngA might be involved in a different glycosylation process or is active under different growth conditions or e.g. during the sporulation process.

In accordance with previous studies (Promadej et al., 1999, Kho and Meredith, 2018), we propose that GtcA acts as a flippase to transport C_55_-P-sugar intermediates across the membrane, which are afterwards used for the glycosylation of LTA in *B. subtilis* and *L. monocytogenes*. Recently, MurJ has been identified as the lipid II flippase involved in peptidoglycan biosynthesis. *In vitro* and *in vivo* studies suggest that MurJ functions by an alternating-access mechanism to transport lipid II across the cell membrane, which is dependent on a membrane potential and potentially also the binding of a cation (Kuk et al., 2019, Kuk et al., 2017, Zheng et al., 2018). MurJ contains 14 transmembrane helices, whereas the putative flippase GtcA only possesses four transmembrane helices. To accommodate the C_55_-P-sugar substrate, we hypothesize that GtcA needs to form a homodimer. GtcA could function similar to MurJ by an alternating-access mechanism. To this end, the GtcA dimer would form an inward-facing cavity, which is bound by the C_55_-P-sugar-intermediate. The binding of the substrate leads to a conformational change, resulting in the flipping of the C_55_-P-sugar intermediate and the formation of an outward-facing cavity. In the next step, the C_55_-P-sugar-intermediate is released and can be used by GT-C type glycosyltransferases, such as the *L. monocytogenes* GtlB or Lmo1079 enzymes, to transfer the sugar onto the LTA and WTA backbone, respectively. A similar mechanism has also been proposed for the SMR efflux transporter EmrE from *E. coli* (Schuldiner, 2009, Fleishman et al., 2006). EmrE is involved in the efflux of a wide range of aromatic cation antibiotics and its activity also depends on the proton motive force (Paulsen et al., 1993, Yerushalmi et al., 1995, Littlejohn et al., 1992, Grinius and Goldberg, 1994). EmrE has also been shown to confer resistance to ethidium bromide and methyl viologen, suggesting a relaxed substrate specificity (Yerushalmi et al., 1996, Schuldiner et al., 2001, Yerushalmi et al., 1995). EmrE is thought to form an antiparallel homodimer in the cell (Muth and Schuldiner, 2000, Chen et al., 2007, Ubarretxena-Belandia et al., 2003), in which the first three transmembrane helices (TMs) of each monomer form the substrate binding chamber. TM4 of each monomer is required for the dimerization of EmrE (Chen et al., 2007). A highly conserved glutamine residue at position 14 in EmrE, which is located in TM1, is important for substrate and proton binding (Muth and Schuldiner, 2000, Yerushalmi and Schuldiner, 2000). In addition, it has recently been shown that the last amino acid residue of EmrE, H110, releases protons upon drug binding and that the C-terminal tail acts like a gate to prevent proton leakage in the absence of a substrate (Thomas et al., 2018). For GtrA-type proteins such as the *L. monocytogenes* GtcA protein, no motifs or amino acids important for their activity have been described thus far. Here, we used an alignment of 1000 GtcA protein sequences to identify conserved amino acids. We found eight highly conserved amino acid residues and by the expression of GtcA variants carrying mutations in these residues in *L. monocytogenes* we identified residues R95 and N132 as essential for the activity of GtcA. According to topology predictions, R95 and N132 are located in TM3 and TM4, respectively (Fig. 6A). But perhaps even more interestingly, we also identified amino acid substitutions in GtcA, which seem to predominately impacted the glycosylation of WTA (F74A) or LTA (K121A). GtcA residues F74 and K121 are predicted to be located in the loops between TM2 and TM3 and TM3 and TM4, respectively (Fig. 6A). Due to the observation that amino acid residue F74 seems to be only important for WTA glycosylation, one might speculate that this residue is important for the recognition of the C_55_-P-GlcNAc intermediate, but dispensable for the recognition of the C_55_-P-galactose intermediate used for LTA glycosylation and *vice versa* for amino acid residue K121.

Taken together, we could show that the GtrA family protein GtcA is involved in the LTA glycosylation process in *L. monocytogenes* and *B. subtilis*. With this, we also revealed that the same small membrane protein and predicted C_55_-P-sugar flippase is required for the LTA and WTA glycosylation process in the *L. monocytogenes* 1/2a serovar strain 10403S. In addition, we identified amino acid residues in the *L. monocytogenes* GtcA protein, which are essential for function and this might help us deceiver the mechanism by which these proteins function. Such protein variants can help us dissect different steps required for protein function such as protein dimerization, substrate binding and release as well as provide information on substrate specificity.

## AUTHOR CONTRIBUTION STATEMENT

**Jeanine Rismondo:** Conceptualization, Funding acquisition, Investigation, Data analysis, Supervision, Visualization, Writing – original draft preparation. **Talal F. M. Haddad:** Investigation, Writing – review & editing. **Yang Shen:** Investigation, Data analysis, Writing – review & editing. **Martin J. Loessner:** Data analysis, Writing – review & editing. **Angelika Gründling:** Conceptualization, Funding acquisition, Data analysis, Supervision, Writing – original draft preparation.

## ACKNOWLEDGEMENTS

This work was funded by the Wellcome Trust grant 210671/Z/18/Z and MRC grant MR/P011071/1 to AG and the German research foundation (DFG) grant RI 2920/1-1 to JR. TFMH was supported by an Imperial College UROP Bursary Award. We thank Dr. Samy Bulous from the Institute of Food, Nutrition and Health, ETH Zürich, for the technical assistance with the UPLC-MS analysis.

