## Supplementary Material for "GtcA is required for LTA glycosylation in *Listeria monocytogenes* serovar 1/2a and *Bacillus subtilis*"

### SUPPLEMENTAL MATERIAL

### SUPPLEMENTAL TABLES

**Table S1: Bacterial strains used in this study**

| Unique ID | Strain name and resistance | Source |
|---|---|---|
| ***Escherichia coli* strains** | | |
| ANG201 | *E. coli* pCN34; AmpR | (Charpentier et al., 2004) |
| ANG1264 | DH5α pKSV7; AmpR | (Smith and Youngman, 1992) |
| ANG2223 | XL1-Blue pKSV7-Δ*lmo2550*; AmpR | (Rismondo et al., 2018) |
| ANG2802 | TG1 pDG1662; AmpR | (Guerout-Fleury et al., 1996) |
| ANG4243 | XL1-Blue pIMK3; KanR | (Monk et al., 2008) |
| ANG4738 | XL1-Blue pKSV7-Δ*gtlB*; AmpR | (Rismondo et al., 2018) |
| ANG4911 | XL1-Blue pKSV7-Δ*gtcA*; AmpR | This study |
| ANG5026 | XL1-Blue pIMK3-*gtcA*; KanR | This study |
| ANG5093 | XL1-Blue pD1662-$P_{ywcC}$-*ywcC-gtcA*; AmpR | This study |
| ANG5182 | XL1-Blue pIMK3-*csbB-yfhO*; KanR | This study |
| ANG5619 | XL1-Blue pKSV7-Δ*lmo0215*; AmpR | This study |
| ANG5620 | XL1-Blue pIMK3-$gtcA_{A65S}$; KanR | This study |
| ANG5621 | XL1-Blue pIMK3-$gtcA_{N69A}$; KanR | This study |
| ANG5622 | XL1-Blue pIMK3-$gtcA_{V73A}$; KanR | This study |
| ANG5623 | XL1-Blue pIMK3-$gtcA_{F74A}$; KanR | This study |
| ANG5624 | XL1-Blue pIMK3-$gtcA_{F91A}$; KanR | This study |
| ANG5625 | XL1-Blue pIMK3-$gtcA_{R95A}$; KanR | This study |
| ANG5626 | XL1-Blue pIMK3-$gtcA_{K121A}$; KanR | This study |
| ANG5627 | XL1-Blue pIMK3-$gtcA_{N132A}$; KanR | This study |
| ANG5628 | XL1-Blue pIMK3-*his-gtcA*; KanR | This study |
| ANG5629 | XL1-Blue pIMK3-*his*-$gtcA_{A65S}$; KanR | This study |
| ANG5630 | XL1-Blue pIMK3-*his*-$gtcA_{N69A}$; KanR | This study |
| ANG5631 | XL1-Blue pIMK3-*his*-$gtcA_{V73A}$; KanR | This study |
| ANG5632 | XL1-Blue pIMK3-*his*-$gtcA_{F74A}$; KanR | This study |
| ANG5633 | XL1-Blue pIMK3-*his*-$gtcA_{F91A}$; KanR | This study |
| ANG5634 | XL1-Blue pIMK3-*his*-$gtcA_{R95A}$; KanR | This study |
| ANG5635 | XL1-Blue pIMK3-*his*-$gtcA_{K121A}$; KanR | This study |
| ANG5636 | XL1-Blue pIMK3-*his*-$gtcA_{N132A}$; KanR | This study |
| ***Bacillus subtilis* strains** | | This study |
| ANG1691 | 168; trpC2 | (Burkholder and Giles, 1947) |
| ANG2749 | 168Δ*csbB::kan*; KanR | (Rismondo et al., 2018) |
| ANG5047 | 168Δ*gtcA::kan*; KanR | This study |
| ANG5102 | 168Δ*gtcA::kan amyE::*$P_{ywcC}$-*ywcC-gtcA*; KanR CamR | This study |
| ***Listeria monocytogenes* strains** | | |
| ANG1263 | 10403S; serovar 1/2a; StrepR | (Bishop and Hinrichs, 1987) |
| ANG2325 | 10403SΔ*gtlA*; StrepR | (Percy et al., 2016) |
| ANG2794 | 10403SΔ*lmo1079*; StrepR | (Rismondo et al., 2018) |

| | | |
|---|---|---|
| ANG4264 | 10403SΔ*gtlB*; StrepR | (Rismondo et al., 2018) |
| ANG4972 | 10403SΔ*gtcA*; StrepR | This study |
| ANG5031 | 10403SΔ*gtcA* pIMK3-*gtcA*; StrepR KanR | This study |
| ANG5195 | 10403SΔ*gtlA*Δ*gtlB*; StrepR | This study |
| ANG5197 | 10403SΔ*lmo1079*Δ*lmo2550*; StrepR | This study |
| ANG5199 | 10403SΔ*gtlA*Δ*gtlB* pIMK3-*csbB-yfhO*; StrepR KanR | This study |
| ANG5203 | 10403SΔ*lmo1079*Δ*lmo2550* pIMK3-*csbB-yfhO*; StrepR KanR | This study |
| ANG5638 | 10403SΔ*lmo0215*; StrepR | This study |
| ANG5639 | 10403SΔ*gtcA* pIMK3-$gtcA_{A65S}$; StrepR KanR | This study |
| ANG5640 | 10403SΔ*gtcA* pIMK3-$gtcA_{N69A}$; StrepR KanR | This study |
| ANG5641 | 10403SΔ*gtcA* pIMK3-$gtcA_{V73A}$; StrepR KanR | This study |
| ANG5642 | 10403SΔ*gtcA* pIMK3-$gtcA_{F74A}$; StrepR KanR | This study |
| ANG5643 | 10403SΔ*gtcA* pIMK3-$gtcA_{F91A}$; StrepR KanR | This study |
| ANG5644 | 10403SΔ*gtcA* pIMK3-$gtcA_{R95A}$; StrepR KanR | This study |
| ANG5645 | 10403SΔ*gtcA* pIMK3-$gtcA_{K121A}$; StrepR KanR | This study |
| ANG5646 | 10403SΔ*gtcA* pIMK3-$gtcA_{N132A}$; StrepR KanR | This study |
| ANG5647 | 10403SΔ*gtcA* pIMK3-*his-gtcA*; StrepR KanR | This study |
| ANG5648 | 10403SΔ*gtcA* pIMK3-*his*-$gtcA_{A65S}$; StrepR KanR | This study |
| ANG5649 | 10403SΔ*gtcA* pIMK3-*his*-$gtcA_{N69A}$; StrepR KanR | This study |
| ANG5650 | 10403SΔ*gtcA* pIMK3-*his*-$gtcA_{V73A}$; StrepR KanR | This study |
| ANG5651 | 10403SΔ*gtcA* pIMK3-*his*-$gtcA_{F74A}$; StrepR KanR | This study |
| ANG5652 | 10403SΔ*gtcA* pIMK3-*his*-$gtcA_{F91A}$; StrepR KanR | This study |
| ANG5653 | 10403SΔ*gtcA* pIMK3-*his*-$gtcA_{R95A}$; StrepR KanR | This study |
| ANG5654 | 10403SΔ*gtcA* pIMK3-*his*-$gtcA_{K121A}$; StrepR KanR | This study |
| ANG5655 | 10403SΔ*gtcA* pIMK3-*his*-$gtcA_{N132A}$; StrepR KanR | This study |

**Table S2: Primers used in this study**

| Number | Name | Sequence |
|---|---|---|
| ANG2979 | gtcA up fw2 | GCGCGGATCCGGTAGTAAGGATAAGACACTGG |
| ANG2980 | gtcA up rev2 | CAAGCAAGATTAGTTCATACTATGTCTTCTTTCTCTC |
| ANG2981 | gtcA down fw2 | CATAGTATGAACTAATCTTGCTTGTTTGCTTCAAC |
| ANG2982 | gtcA down rev2 | GCGCGGTACCGATCATCCAGTACATCAGACGG |
| ANG3036 | gtcA pIMK3 fw NcoI | CATGCCATGGGGAACAAAATAAGAAAATGGTTAGACAAG |
| ANG3037 | gtcA pIMK3 rev SalI | GCGCGTCGACTTATTTTTCACTTTGAAAATGATCC |
| ANG3068 | fw 1kb gtcA | CACATAGTTGCTCGGTGTTC |
| ANG3069 | Front ApaI gtcA | CGCGGGCCCGGTTGTGAAAACCCCCATAATG |
| ANG3070 | Back XhoI gtcA | GCCGCTCGAGTTTAAAAAAACAAAAGAAGAAGGGG |
| ANG3071 | rev 1 kb gtcA | GCCGTCTTAATAGCACGATG |
| ANG3089 | 5-BamHI-ywcC with P | CGCGGATCCAGTTTTCATAGTAACCTCCAATTG |
| ANG3090 | 3-HindIII-gtcA BSU | CCCAAGCTTGCTGCAGTCCTCTCATACC |
| ANG3143 | pIMK3-*csbB-yfhO* fw | GCGCCCATGGGGAAGCAAGGATTAATCTCG |
| ANG3144 | pIMK3-*csbB-yfhO* rev | GCGCGTCGACCTATATAGAGCCGGGCTTTTTAC |
| ANG3215 | gtcA A65S fw | CGTCCTTTTTTCATATTTTTCTAATAAAAAATATGTTTTTG |
| ANG3216 | gtcA A65S rev | TAGAAAAATATGAAAAAAGGACGGATGCAACC |
| ANG3217 | gtcA N69A fw | CATATTTTTCTGCTAAAAAATATGTTTTTGAAAGCTATAC |
| ANG3218 | gtcA N69A rev | CATATTTTTTAGCAGAAAAATATGCAAAAAGGACGG |
| ANG3219 | gtcA V73A fw | TAAAAAATATGCTTTTGAAAGCTATACACCTACTTG |
| ANG3220 | gtcA V73A rev | AGCTTTCAAAAGCATATTTTTTATTAGAAAAATATGCAAAAAG |
| ANG3221 | gtcA F74A fw | AAAATATGTTGCTGAAAGCTATACACCTACTTGG |
| ANG3222 | gtcA F74A rev | GTATAGCTTTCAGCAACATATTTTTTATTAGAAAAATATGCAA |
| ANG3223 | gtcA F91A fw | GTTACATCAGCTTTCGGTTTCCGCTTTTTGAC |
| ANG3224 | gtcA F91A rev | GGAAACCGAAAGCTGATGTAACTTCTCGAGCGC |
| ANG3225 | gtcA R95A fw | TTTCGGTTTCGCCTTTTTGACTTATTTAGTGGACATTT |
| ANG3226 | gtcA R95A rev | AGTCAAAAAGGCGAAACCGAAAAATGATGTAACTTC |
| ANG3227 | gtcA K121A fw | ATTATGGGCAGCAATTTGGACGAATGTAATTGTCC |
| ANG3228 | gtcA K121A rev | TCGTCCAAATTGCTGCCCATAATTCATTTATAGATAAAA |
| ANG3229 | gtcA N132A fw | CCTTGTACTTGCTTATGTGTTTAGTAAATGGATCATTT |
| ANG3230 | gtcA N132A rev | CTAAACACATAAGCAAGTACAAGGACAATTACATTCG |
| ANG3305 | lmo0215 up fw BamHI | CGCGGATCCCTGCGATGGTTCCTGATG |
| ANG3306 | lmo0215 up rev | TAATTTATATCTCATTAGTCGCTTCATGGAACGTTCTGACAC |
| ANG3307 | lmo0215 down fw | AAGCGACTAATGAGATATAAATTACTCGGTCCAAAAGATTTTG |
| ANG3308 | lmo0215 down rev KpnI | CGGGGTACCCTCGGCTCTGACTTACTC |
| ANG3345 | His-*gtcA* fw 2 | CACCACCACCACAACAAAATAAGAAAATGGTTAGAC |
| ANG3346 | NcoI-His 2 | CATGCCATGGGGCACCACCACCACCACCACAACAAA |

**SUPPLEMENTAL FIGURES AND LEGENDS**

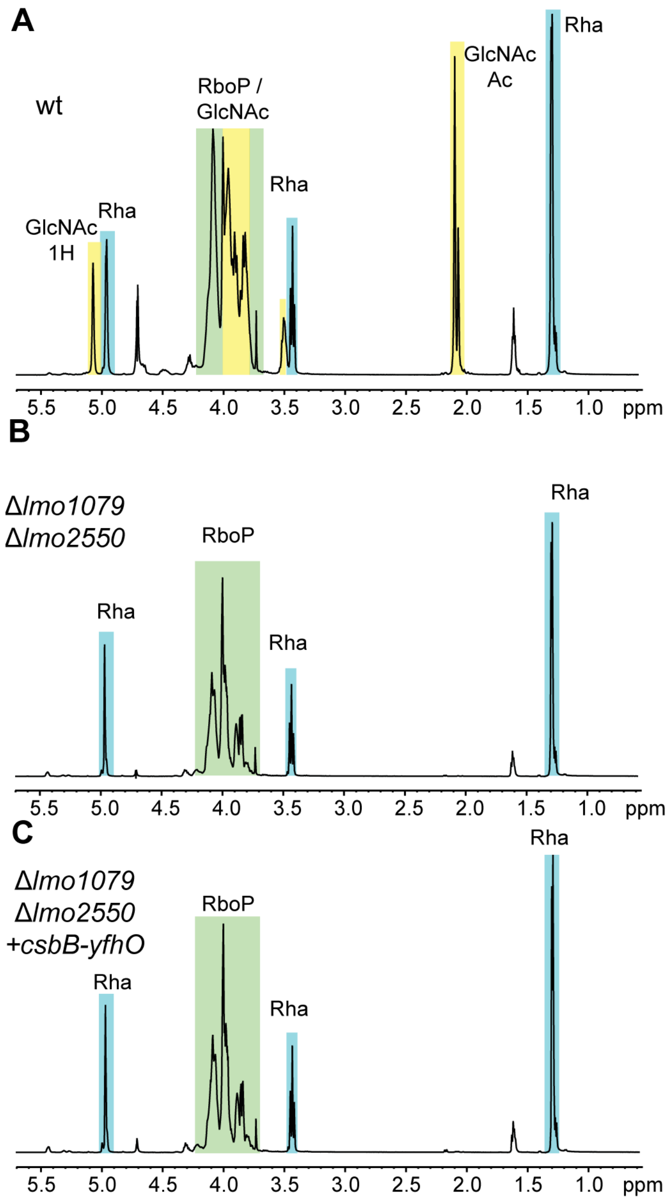


**Figure S1: NMR analysis of WTA isolated from wildtype *L. monocytogenes* strain 10403S, the *lmo1079/lmo2550* mutant and the *lmo1079*/*lmo2550* mutant expressing the *B. subtilis csbB-yfhO* operon.** (A-C) NMR spectra of WTA. WTA was extracted from *L. monocytogenes* (A) 10403S (wt), (B) the 10403SΔ*lmo1079*/Δ*lmo2550* mutant and (C) strain 10403SΔ*lmo1079* Δ*lmo2550*+*csbB-yfhO* (grown in presence of IPTG) and analyzed by NMR. Peaks of nonexchangeable protons were assigned to the different WTA components using previously published spectra and highlighted in colored boxes (Kamisango et al., 1983, Reichmann et al., 2013, Brauge et al., 2016). One representative result of two independent experiments is shown.

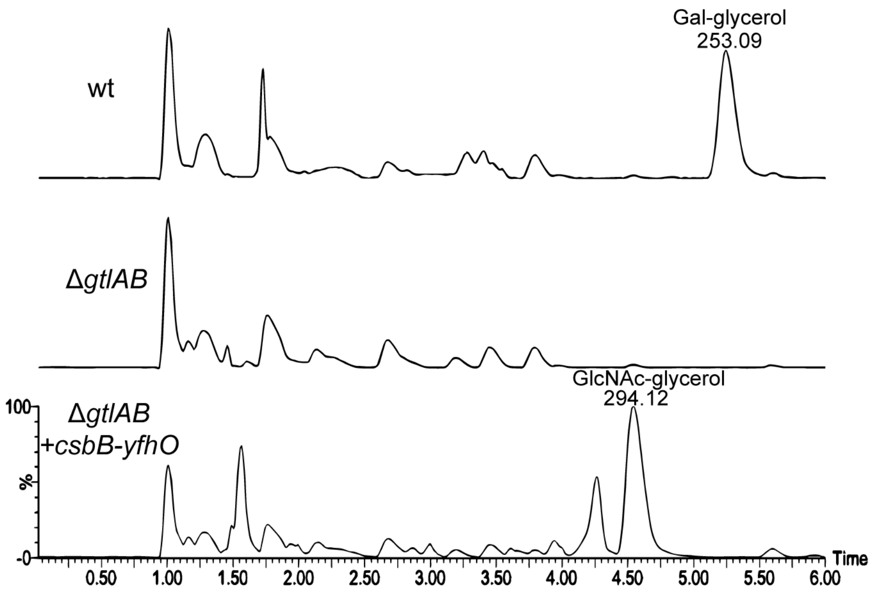


**Figure S2: UPLC-MS/MS analysis of LTA fragments isolated from wildtype *L. monocytogenes* 10403S, the *gtlAB* mutant and the *gtlAB* mutant expressing the *B. subtilis csbB-yfhO* operon.** Purified LTA was analyzed by UPLC-MS/MS as described in the methods section. Peaks corresponding to Gal-glycerol with a m/z of 253.09 and GlcNAc-glycerol with a m/z of 294.12 are labelled. The highest peak for the LTA sample extracted from strain 10403SΔ*gtlAB+csbB-yfhO* was set to 100%. One representative result of three independent experiments is shown.

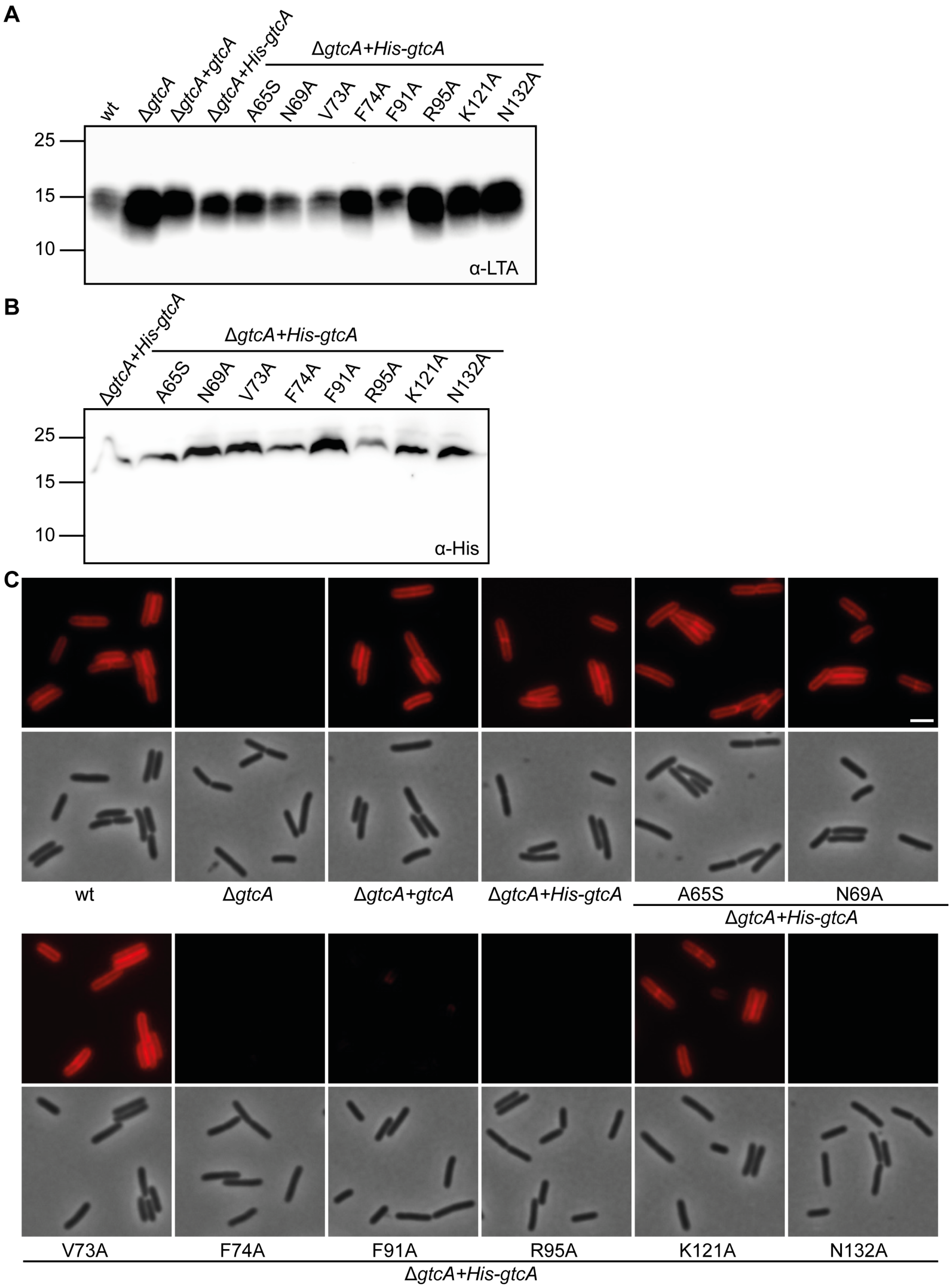


**Figure S3: Identification of amino acid residues essential for the function and/or stability of His-tagged versions of the *L. monocytogenes* 10403S GtcA protein.** (A) Detection of LTA by western blot. Cell extracts of the indicated *L. monocytogenes* strains expressing His-GtcA or derivatives thereof were prepared, separated on a 15% SDS PAGE and LTA detected using a polyglycerolphosphate-specific monoclonal antibody. (B) Detection of His-GtcA proteins by western-blot. Protein extracts of the indicated *L. monocytogenes* strains were prepared as described in the method section and separated on a 15% Tricine SDS PAGE gel. His-GtcA proteins were detected using a monoclonal poly-Histidine-Peroxidase antibody.

(C) Microscopy analysis and detection of WTA glycosylation using the fluorescently labelled WGA-Alexa 594 lectin. Log-phase cells of the indicated *L. monocytogenes* strains were stained with WGA-Alexa 594 as described in the methods section and subjected to phase contrast and fluorescence microscopy. Scale bar is 2 µm. One representative result of three independent experiments is shown.

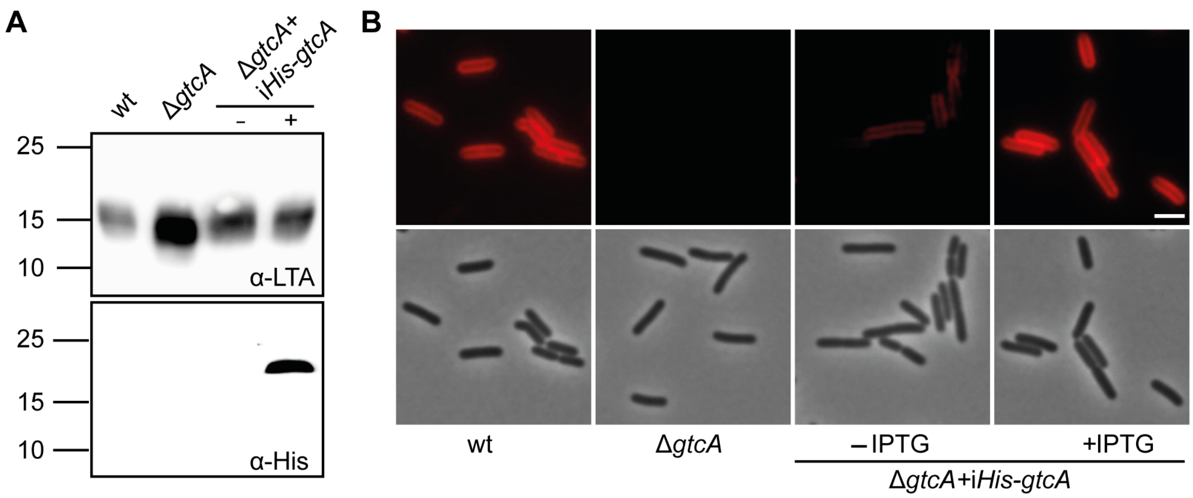


**Figure S4: Complementation analysis using *L. monocytogenes* serovar 1/2a strain 10403SΔ*gtcA* expressing His-tagged GtcA.** (A) Detection of LTA and His-GtcA by western blot. Cell and protein extracts of *L. monocytogenes* strains 10403S, 10403SΔ*gtcA* and strain 10403SΔ*gtcA* pIMK3-*His-gtcA* grown in the absence or presence of IPTG were prepared as described in the methods section. Upper panel: Cell extracts were loaded on a 15% SDS PAGE gel and LTA was detected by western blot using a polyglycerol-phosphate-specific monoclonal antibody. Lower panel: Protein extracts were loaded on 15% Tricine SDS PAGE gels and the His-GtcA protein were detected using a monoclonal polyHistidine-Peroxidase antibody. (B) Microscopy analysis and detection of WTA glycosylation using the lectin WGA-Alexa 594. Log-phase cells of the indicated *L. monocytogenes* strains were stained with the WGA-Alexa 594 dye as described in the methods section and subjected to phase and fluorescence microscopy. Scale bar is 2 µm. One representative result of three independent experiments is shown.
